# The Environmental Games

**DOI:** 10.64898/2026.07.18.739357

**Authors:** Andrew Marcus

## Abstract

For over 4 billion years, the environment has guided evolutionary selection by presenting biological entities with opportunities and challenges and by mediating their interactions. In evolutionary game theory (EGT), the replicator equation defines the game according to direct interactions between biological entities. Eco-evolutionary models have described environmental feedback by modeling environmental states dynamically. Yet, these models channel environmental pressures through the direct interaction coefficients instead of creating a distinct game. Here, I invent Pure Environmental Games (PEGs) and formally introduce the environment as a strategy in EGT. The PEGs describe the neutral selection game and environmental games operating in parallel. Through the net specific accumulation rates of microbes parametrized by the environmental state, Environmental Biotechnology—an engineering discipline designing wastewater treatment plants worldwide — offers a collection of instances of PEGs. The environmental strategy resists invasion from biological strategies when it is a strict Nash equilibrium. The PEGs provide a template for identifying regimes of environmental selection within an environmental game (e.g., the r/K selection game). This mapping identifies persistence as a new dynamic regime under extinction. In a closed system, life persists by partnering with the environmental strategy consistent with Schrödinger’s negative entropy.

## Introduction

The environment is the “invisible” hand directing evolution. As early lifeforms invaded uninhabited frontiers, the environment selected its new residents by presenting both opportunities and challenges. Yet, the environment has not been formally introduced as a strategy in evolutionary game theory (EGT).

The replicator equation (Figure 1a) and Lotka-Volterra model (LVM) (Figure 1b) are foundational equations in the EGT and population ecology, respectively. Hofbauer and Sigmund ^1^ proved the equivalence of the two models with *n* LVM species mapping onto *n* + 1 replicator strategies. This mapping separates the intrinsic growth rates under *S*_0_ from the direct interaction coefficients in the payoff matrix (*S*_*n*_ for *n* ≥ 1, Figure 1d). Traditionally, the direct interaction coefficients define the game^2^. When the intrinsic growth rates are assumed to be identical across different strategies, traditional EGT classifications map to replicator dynamics (Supplementary Note S2;^3^).

**Figure 1.**
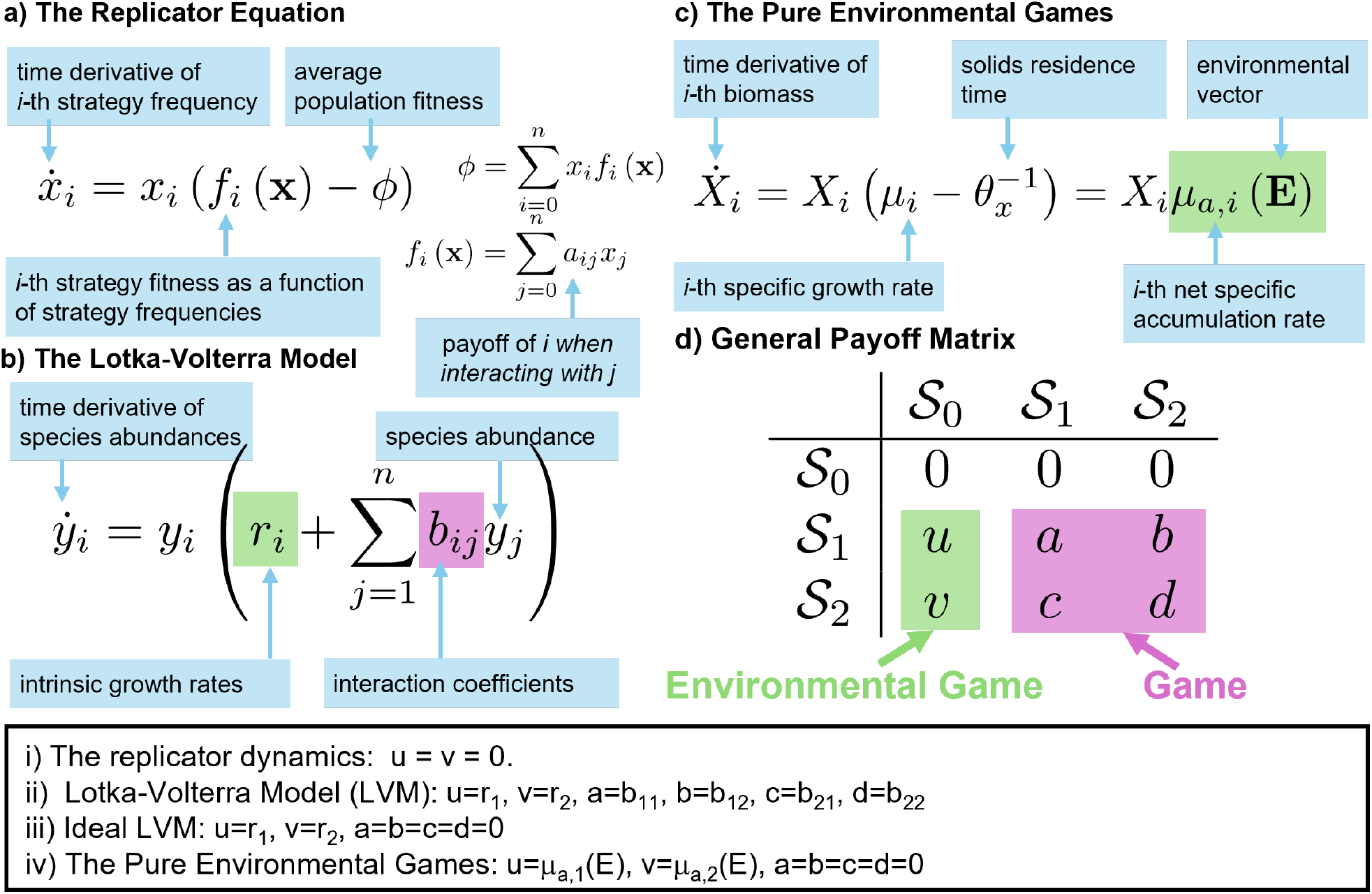
a) The replicator equation models the rate of change in strategy frequency 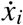, where the subscript *i* indexes strategies. The frequency increases when its fitness, *f*_*i*_, exceeds the average population fitness, *ϕ. f*_*i*_ is determined by summing the product of the payoff coefficient, *a*_*i j*_, by *x* _*j*_ . *ϕ* is determined by summing the products of *f*_*i*_ and *x*_*i*_. b) The Lotka-Volterra model (LVM) models the rate of change in species abundance 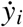. The LVM has two parameters: intrinsic growth rate, *r*_*i*_, and the interaction coefficient, *b*_*i j*_, c) The Pure Environmental Games (PEGs) model the biomass concentration in the absolute coordinate 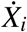. The PEGs have one coefficient, the net specific accumulation rate, *μ*_*a,i*_, which is parameterized by the environmental vector, **E**. d) The general payoff matrix summarizes the relationship between three equations for the case of two biological strategies. The replicator equation models the competition of biological strategies. The first row is set to zero (i.e., *f*_0_ = 0) to allow the replicator equation to track the total abundance or concentration of biological components in the absolute coordinate. The coefficients *a, b, c*, and *d* encode the payoff resulting from direct interactions between strategies. The LVM models the ecological interactions between species in the absolute coordinate. When the LVM is mapped to the replicator equation according to Hofbauer and Sigmund^1^, the coefficients *u* and *v* equal the intrinsic growth rates. The ideal LVM can be derived by setting *a* = *b* = *c* = *d* = 0. The PEGs model systems driven by interactions of biological strategies with the environmental strategy. Mass conservation relationships from Rittmann and McCarty^13^ provide abundant instances of PEGs. Structurally, the PEGs are analogous to the ideal LVM, and *u* and *v* equal the net specific biomass accumulation rates, *μ*_*a,i*_ (**E**).

Among EGT classifications, neutral selection corresponds to a game where deterministic replicator dynamics no longer control biological behavior (Figure 2b; Chapter 4 of Nowak ^3^ ). However, when environmental conditions differentiate the intrinsic growth rates of biological entities^2,4^, the intrinsic growth rates can drive selection even in the absence of replicator dynamics^2^. A differential in intrinsic growth rates alters the selection baseline against which direct interspecies interactions operate; thereby, mapping between the game and EGT classifications is disrupted^2^. Therefore, Tarnita and Traulsen ^2^ call for the field of EGT to embrace approaches that incorporate a more nuanced ecology, such as Lotka-Volterra or adaptive dynamics.

**Figure 2.**
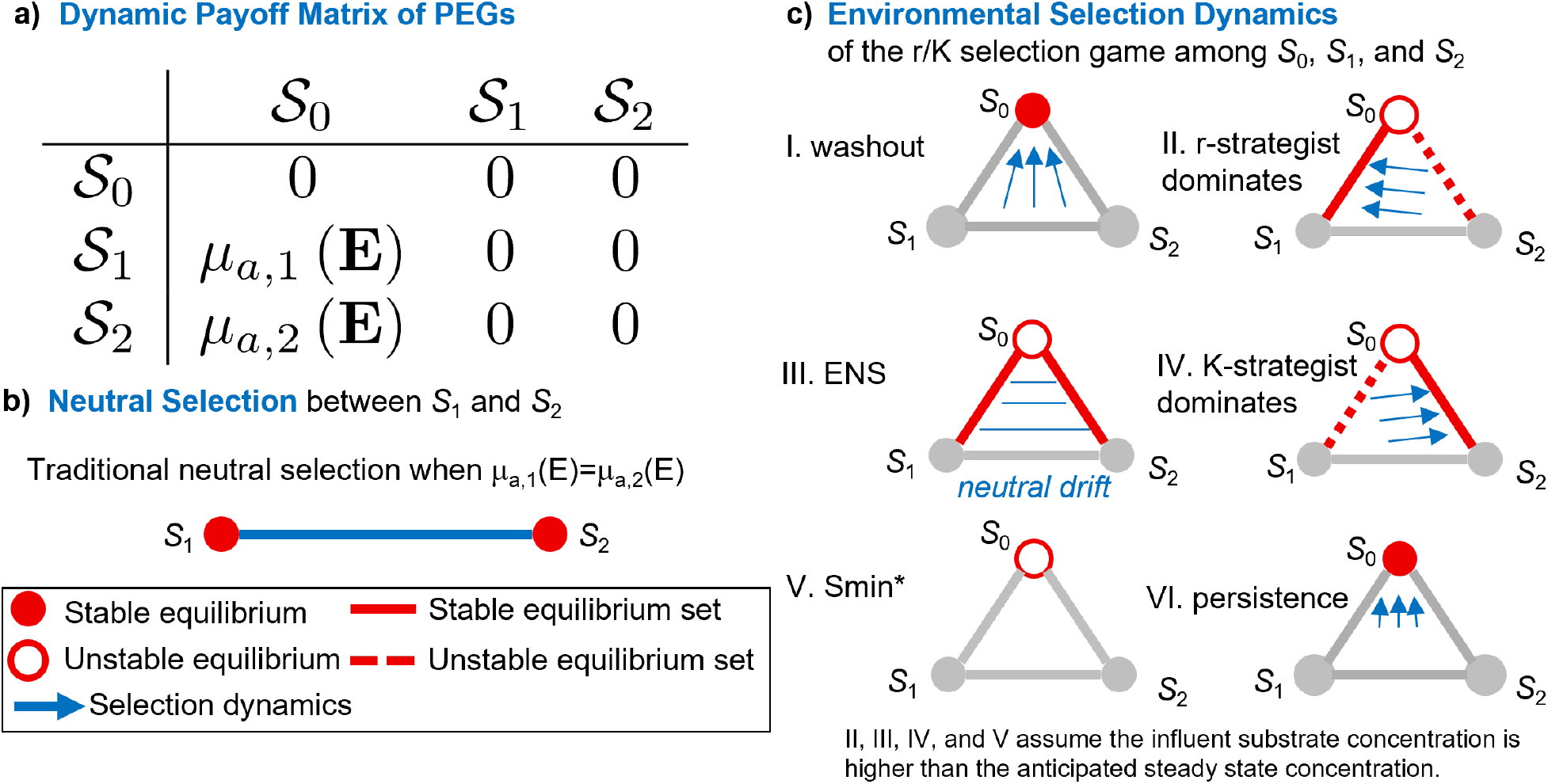
a) The dynamic payoff matrix for the Pure Environmental Games (PEGs). b) The PEGs demonstrate the traditional neutral selection dynamics when *μ*_*a*,1_ (**E**) = *μ*_*a*,2_ (**E**). c) Six environmental regimes of the r/K environmental game that operate in parallel to neutral selection. I. In washout 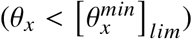, biological strategies cannot grow fast enough and cannot invade the environmental strategy. II. The r-strategist dominates by establishing a stable dynamic equilibrium. III. At 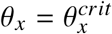, the anticipated steady state substrate concentrations 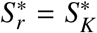 (i.e., environmentally neutral selection, ENS). Thus, the r-strategist and K-strategist are environmentally neutral. IV. The K-strategist dominates. V. Under 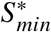, infinitesimally small biomass establishes a stable dynamic equilibrium. VI. In the extinction regime, persistence becomes an evolutionary strategy.

Past eco-evolutionary models have dynamically tracked absolute quantities (such as population density and environmental state) and modified direct interaction coefficients to describe the environmental feedback^5–8^. While this approach has the potential to bring nuanced ecology, it does not dissolve the difficulty of mapping the game to EGT classification when the intrinsic rate differentials exist.

Here, in response, I invent a new evolutionary game beyond the direct interaction coefficients in two steps. Step 1 derives an ideal LVM framework by setting the direct-interaction submatrix to null. Step 2 sets the payoffs that biological strategies derive from the environmental strategy to the net specific accumulation rates parameterized by the environmental state, yielding the Pure Environmental Games (PEGs). Thereby, the PEGs will provide a more ecologically nuanced interpretation of the neutral selection game by visiting the evolutionarily stable strategy (ESS) classifications and the r/K selection game as an instance of PEGs. While the main text provides high-level ideas, the Supplementary Note provides the following domain-specific concepts and derivations for a broad readership:

- Note 1 provides a technical overview.
- Notes 2–3 establish EGT foundations and the replicator equation and Lotka-Volterra equivalence.
- Note 4 derives the ideal LVM, which serves as a skeleton and a precursor to the PEGs.
- Note 5 defines the PEGs and identifies Environmental Biotechnology as a collection of instances of PEGs to import environmental selection principles and conservation laws.
- Note 6 dynamically models environmental selection operating in parallel to the limiting case of neutral selection using *Methanosarcina*-*Methanothrix* as a canonical example.

This work aims to serve as a foundation for cross-disciplinary conversations between EGT, Environmental Biotechnology, and Microbial Ecology — and other curious minds — rather than to claim novel predictions.

### Pure Environmental Games

Imagine biological entities as gas molecules. At low temperatures or under high density, gas molecules interact and behave non-ideally. At high temperatures or under low density, the interactions become negligible, revealing the ideal gas law. In a similar spirit, removing interactions between biological entities “purifies” environmental selection.

Begin by setting the game to the limiting case of neutral selection where *a* = *b* = *c* = *d* = 0 (Figure 2a). This produces the ideal LVM, where deterministic selection can continue via the intrinsic growth rates (*u* and *v* in Figure 1d). Equating the intrinsic growth rates to the payoffs that biological strategies receive from the environmental strategy, *μ*_*a,i*_ (**E**), produces the PEGs (Figures 1c and 2a).

The general payoff matrix shows the relationship between PEGs and traditional EGT (Figure 1d) in two parts: the direct interaction submatrix defines the game in traditional EGT; the intrinsic growth submatrix defines the environmental game. By setting the game to null, the PEGs describe the neutral selection game operating in parallel to the environmental game. The PEGs depart from the traditional EGT by making *S*_0_ the environmental strategy and *x*_0_ the environmental frequency. The asymmetry between one environmental strategy and *n* biological strategies indicates that all biological strategies share the same local environment. Multiple environmental strategies may exist in the presence of environmental variation, though spatial games are beyond the scope of this work. As with the traditional EGT, the biological strategies still describe a fixed or “hardwired” phenotype. The payoffs can change, but the underlying representation does not. Thereby, *μ*_*a,i*_ (**E**) provides a mathematical means for the environmental pressure to alter the fitness of *i*-th biological entity when multiplied by *x*_0_.

By placing *μ*_*a,i*_ (**E**) in the general payoff matrix, the environmental strategy can be interpreted through the evolutionary stability concepts with a caveat. Static equilibrium concepts describe strategies in relation to the payoff structure. Because the environmental state is a dynamic variable, the payoff matrix becomes dynamic. Consequently, the ESS classification of a strategy (e.g., *S*_0_) becomes state dependent.

### Evolutionarily Stable Strategy

The concept of ESS was introduced by Smith and Price ^9^ as a static equilibrium to describe strategies that, if adopted by the majority, resist invasions by rare mutants. Consider a PEG with one biological strategy with the following payoff structure, equation (1):

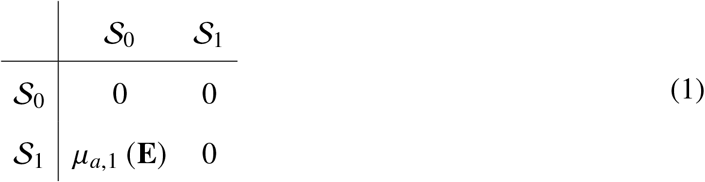

On the one hand, the biological strategy, *S*_1_, is always a Nash equilibrium, and its ability to invade depends entirely on the environmental state. On the other hand, *S*_0_ is a strict Nash equilibrium and resists invasion if *μ*_*a*,1_ (**E**) < 0. *S*_0_ disqualifies as a (strict) Nash equilibrium when *μ*_*a*,1_ (**E**) > 0 and the biological strategy can invade the environment. At a steady state, the payoff matrix becomes null, and the environmental strategy is a Nash equilibrium (not strict). These ESS classifications are independent of the implementation of *μ*_*a*,1_ (**E**).

It should be noted that the ESS classification is based on static equilibrium concepts, and a null payoff matrix does not imply the absence of selection dynamics. The selection dynamics will depend on the implementation of *μ*_*a,i*_ (**E**) and parameterization of the environmental game. The PEGs take a modular approach, and *μ*_*a,i*_ (**E**) can be set to any specific accumulation rate that depends on the environmental state. Environmental Biotechnology provides a collection of *μ*_*a,i*_ (**E**) implementations as biomass balances. The next section presents one such instance, the classic r/K game, and describes its environmental selection dynamics.

### Environmental Selection Dynamics

Environmental Biotechnology is an academic discipline rooted in sanitary engineering – one of the defining public health achievements of the 20th century. Environmental Biotechnology brings to PEGs rich principles of environmental selection, integrated into biological models, empirically validated through the design, operation, and control of tens of thousands of biological wastewater treatment processes worldwide^10–13^. The solids residence time, *θ*_*x*_, is a fundamental parameter for environmental selection^13^. In practice, engineers can set *θ*_*x*_ to enrich for stable microbiological communities with desirable attributes. In a medical context, *θ*_*x*_ may be defined by physiology, such as in the human digestive system^14^. This section identifies six dynamic regimes within the r/K selection game operating in parallel to the neutral selection game with *Methanosarcina* and *Methanothrix* as a canonical example of environmental succession. The analysis assumes an idealized chemostat (spatially homogeneous and no wall effects) (Supplementary Note S6). For simplicity, the energy dissipated by microorganisms is assumed to be transported out of the system by the bulk fluid (Supplementary Note S5). Parameters for *Methanosarcina* and *Methanothrix* are well-grounded in experimental data, microbial kinetics, and thermodynamics^15–17^. Six selection dynamics are identified with Roman numerals (Regimes I-VI) and illustrated conceptually (Figure 2c), which are then identified within the parameter space of *θ*_*x*_ and the influent substrate concentration (*S*^0^) (Figure 3) with full dynamics presented in Supplementary Note S6 (Extended Data Fig. 1-4).

**Figure 3.**
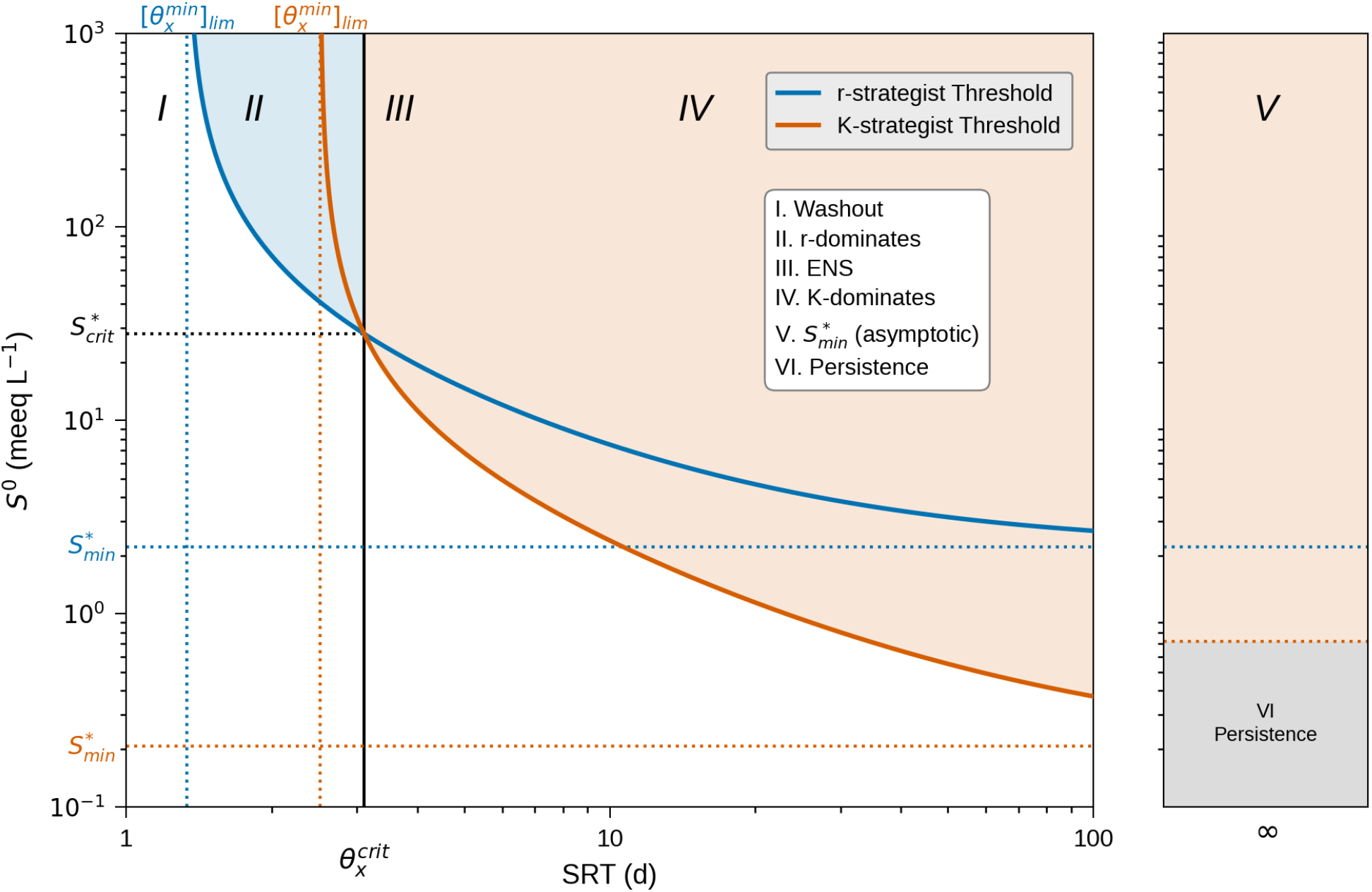
Six environmental selection regimes (Roman numerals I-VI as defined in legend) within the parameter space of the r/K environmental game. x-axis and y-axis show the solids residence time (*θ*_*x*_, d) and influent substrate concentration (*S*^0^; meeq L−1), respectively. Blue and Vermillion lines describe the influent substrate thresholds for establishing steady state biomass. They are obtained by calculating the steady state substrate concentration for r-strategist (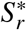; meeq L−1) and K-strategist (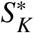; meeq L−1), respectively. The regions above 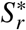 and 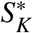 are color-filled to identify where respective strategists dominate: blue – r-strategist dominates and red – K-strategist dominates. The left panel and right panels describe finite *θ*_*x*_ and the asymptotic limit as *θ*_*x*_ approaches infinity, respectively. Dotted lines describe asymptotic behaviors around 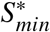 and 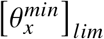 as labelled. For *Methanothrix*, Regimes I-IV used *K*_*Ac*_ = 6.3 meeq L−1 and Regimes V-VI *K*_*Ac*_ = 22 meeq L−1 as indicated in Extended Data Table 2.

### Regime I) Washout

Monod kinetics defines the minimum solids residence time required to prevent washout, 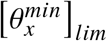 for each type of microbe^18^ (Extended Data Fig. 1). When 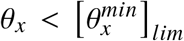, *μ*_*a,i*_ (**E**) < 0 for all biological strategies, the environmental strategy is a strict Nash equilibrium. The system removes microbes faster than they can grow, and the environmental strategy resists invasions. Although 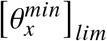 sets a strict condition for washout, the white region below the curve in Figure 3 (*S*^0^ < *S*^*^) also is generally known as washout.

### Regimes II and IV) r/K Selection

Microbes require sufficient time and energy to invade the environmental strategy. When 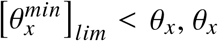 sets the anticipated steady-state substrate concentration, *S*^*^ ^18^. If *S*^0^ > *S*^*^, *μ*_*a,i*_ (**E**) > 0. Consequently, the environmental strategy disqualifies as a (strict) Nash equilibrium. Microorganisms can invade the environment, and the environment can, in turn, select biological strategies.

r/K selection is a classic microbial ecology model for ecological succession. The r-strategist represents opportunists who thrive in volatile environments (*Methanosarcina*). The K-strategist represents gleaners thriving in stable environments (*Methanothrix*). For describing competitive exclusion, *S*^*^ is the same quantity as Tilman’s *R*^*^ derived for different systems^18,19^. Those microbes with a lower *S*^*^ can completely exclude their competitors^20^ (Supplementary Note S5 and S6). The critical solids residence time, 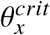 marks the transition between the selection dynamics. When 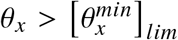 and 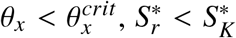 and the r-strategists can invade the environmental strategy to establish a steady presence and dominate over K-strategists (Extended Data Fig. 2). Similarly, K-strategists dominate over r-strategists when 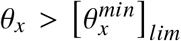 and 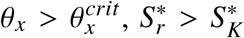 (Extended Data Fig. 3).

### Regime III) Environmentally Neutral Selection

Between Regimes II and IV, environmental neutrality occurs at 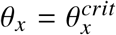, which ensures 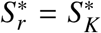 and *μ*_*a*,1_ (**E**) = *μ*_*a*,2_ (**E**) at steady state (see Supplementary Note S6). As with neutral selection in replicator dynamics^3^, deterministic environmental selection dynamics no longer describe the outcome of r/K selection. In principle, exclusion of microbes with identical environmental fitness can be described by neutral drift^3,21^. In practice, however, rand K-strategists such as *Methanosarcina* and *Methanothrix* do coexist in biological reactors due to non-ideal conditions, and their coexistence can be promoted through mechanisms such as feeding controls, cross-feeding, and spatial heterogeneity^15^.

### Regime V) 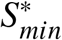

As *θ*_*x*_ approaches infinity, the steady-state substrate concentration asymptotically approaches 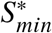, which represents the minimum substrate concentration that can maintain steady-state biomass^22^. At 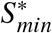, the minimum energy dissipation rate for sustaining a steady-state biomass for *Methanothrix* is 1.4 × 10−11 kJ cell−1 y−1 (Extended Data Fig. 4).

### Regime VI) Persistence

Below 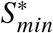, microorganisms decay exponentially, which is described mathematically as fast extinction^23^. However, extinction can be slow enough to be relevant in terms of physical time, as microorganisms can shut down their metabolism to the bare minimum of maintenance requirements^24^. Deep-ocean sediment is among the few natural systems known to approximate an isolated system. Deep-ocean microorganisms may persist for thousands or millions of years by reducing the energy dissipation rate to as low as 10−15 kJ cell−1 y−1 ^24^, a 4-order-of-magnitude reduction compared to *Methanothrix* at 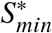. Persistence is relevant in environmental contexts where microorganisms buy time until invasion becomes possible, using mechanisms such as metabolic dormancy and sporulation.

### Significance

Describing the immense diversity of this planet, represented largely by microorganisms, requires an evolutionary language^25,26^. Earth provides life with countless environmental variations, which serve as the foundation for building immense biodiversity. Therefore, a tool is needed to ask how life invades the environment.

In traditional EGT, the direct interaction submatrix defines the game^2^. Eco-evolutionary models have incorporated environmental feedback into the direct interaction matrix^5–8^. The PEGs depart from the tradition by creating the environmental game that operates in parallel to the neutral selection game. All environmental pressures were ensured to enter through the environmental game by setting the direct interaction submatrix (the game) to null (the limiting case of neutral selection).

The PEGs formally introduce the environment as a strategy. By placing the net accumulation rates of microorganisms in the environmental game submatrix, the PEGs recruit over a century of accumulated knowledge of environmental selection from Environmental Biotechnology into evolutionary games. r/K selection was chosen as an instance of an evolutionary game because its microbial kinetics parameters can be estimated from first principles^17,27^ and are established empirically^16^. The r/K selection game demonstrates its environmental selection is bound by time 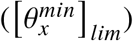 and energy (set at 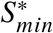)^18,22^. Within this boundary, biological strategies invade the environment and compete, not through direct interactions, but through indirect interactions mediated by the environmental strategy. Having deterministic selection continue under the neutral selection game is consistent with Tarnita and Traulsen^2^ under *r*_1_ ≠ *r*_2_. Deterministic dynamics no longer determine a unique outcome, and neutral drift can occur when *μ*_*a*,1_ (**E**) = *μ*_*a*,2_ (**E**).

By dynamically modeling the environmental state, the PEGs bring empirically grounded knowledge of microbial kinetics and thermodynamics into the EGT. The principles of environmental selection and ESS work elegantly together: thermodynamic feasibility has historically served as a guide for anticipating novel metabolism (e.g., anaerobic ammonium oxidation was anticipated two decades before its discovery^28,29^), and now ESS concepts can define when these novel biological strategies can invade the environmental strategy.

Critically, the PEGs make the system boundary explicit. The PEGs describe mass exchange between the system and the surroundings using *θ*_*x*_, which is also a fundamental parameter in spatially heterogeneous systems such as biofilms. According to Erwin Schrödinger, life feeds on negative entropy to maintain order^30^. In a closed system, established when *θ*_*x*_ reaches infinity, exporting entropy to the surroundings is prohibited. However, deep-sea sediment microorganisms exemplify persistence by dissipating energy through interactions with the environmental strategy.

The PEGs follow a modular design principle. Analyzing the dynamic regimes within an instance of PEGs brings physical nuance to exploring traditional EGT concepts defined mathematically. The persistence regime is particularly important in microbial ecology, where extinction is not as readily observable as in macroecology. Persistence may help to explain resurgences of measles31 and resilience of antibiotic resistance genes^32^. Other instances of environmental games may be considered by modifying the payoff that biological entities receive from the environment.

## Methods

### Conservation Laws

Simulations for environmental selection dynamics were performed by solving mass and energy conservation laws. A general formula for writing chemical reaction is 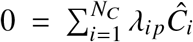 where *Ĉ*_*i*_ is the formula for component *i* [M], *λ*_*i,p*_ is the stoichiometry coefficient for component *i* in reaction *p* [*M*_*i*_ per reaction], and *N*_*C*_ is the number of components. *λ*_*i,p*_ is negative for the reactant and positive for the product. Then, a general mass conservation law can be written as 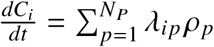 (**E**) where *ρ*_*p*_ is the rate of *p*-th process and *N*_*P*_ is the number of processes. For *Methanosarcina*-*Methanothrix* system, Extended Data Table 1 summarizes the microbiological and transport processes in the standard Gujer format. Extended Data Table 2 summarizes the microbial kinetic parameters.

During growth and decay, microorganisms dissipate energy through anabolic and catabolic reactions. The reaction stoichiometry for each reaction is written as 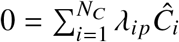 where *Ĉ*_*i*_ is the formula for component *i* [M] and *λ*_*i,p*_ is the stoichiometry coefficient for component *i* in a process *p* [*M*_*i*_ per reaction] when the process *p* is a reaction. The Gibbs free energy of reaction is written as 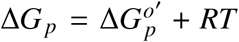 ln 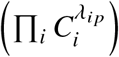 where 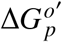 is the standard Gibbs free energy of the reaction [*E M*^−1^], *R* is the ideal gas constant (8.314 J/K-mol), and *T* = 298.15*K*. The superscript “o” denotes standard condition and the prime denotes pH=7. Extended Data Table 3 summarizes the anabolic, catabolic, and endogenous decay reactions for acetoclastic methanogens in terms of eeq transferred (same for *Methanosarcina* and *Methanothrix*). Finally, the specific energy dissipation rate for the *i*-th microbe is calculated as 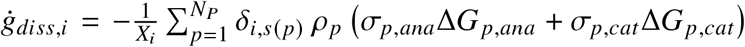. where *X*_*i*_ is the biomass concentration for the *i*-th microbe, *σ*_*p,ana*_ = 1 and *σ*_*p,cat*_ = (1 − *Y*_*i*_)/*Y*_*i*_ are the partitioning of the electron equivalents between anabolic and catabolic reactions per eeq_*X*_ of biomass generated, and *ρ*_*p*_ is the process rate of the *p*-th process. Δ*G*_*p,ana*_ and Δ*G*_*p,cat*_ are the Gibbs free energy of reaction for process *p* (kJ per eeq_*rxn*_) for the reaction defined in Extended Data Table 3. *δ*_*i,s*(*p*)_ defines the Kronecker delta function (*δ*_*i,s*(*p*)_=1 when *i* = *s*( *p*) and 0 otherwise). *s*( *p*) maps the process *p* to its associated biomass type (cin or thr), as given by the rows of the energy dissipation matrix summarized in Extended Data Table 4.

### Numerical Methods

The Lotka-Volterra model (LVM) and the replicator equation are mathematically equivalent^1^. All models, including LVM and conservation laws, were solved in absolute coordinates using `solve_ivp` from Scipy in Python. The consistency of the numerical methods was verified by solving the replicator equations directly and comparing trajectories, yielding a maximum relative error of 3.4 × 10−4. Since the simplex dynamics occur in the transformed time coordinate (*τ*), we solve the following relationships along with the ordinary differential equations: *dτ*/*dt* = 1/*x*_0_.

## Code Availability

All code used to generate the results and figures in this study is available at https://github.com/amarcus1/environmental-games/tree/v1.0. A permanent, versioned archive with a DOI will be deposited via Zenodo upon acceptance and provided at that stage.

## Acknowledgements

This work is dedicated to the memory of my grandmother. The author thanks Drs. John Norton and Dienye Tolofari for creating an intellectually stimulating environment.

## Author Contributions

A.M. is the sole author and is responsible for all aspects of the work.

## Supplementary Information

**Extended Data Table 1:** Gujer matrix for the two-species acetoclastic methanogenesis chemostat model. *X*_*cin*_ is *Methanosarcina* biomass, *X*_*thr*_ is *Methanothrix* biomass, and *S*_*Ac*_ is the acetate concentration. Model parameters are defined in Extended Data Table 2

| Process | Components | | | Process Rate<br>$\rho$ (E) |
| --- | --- | --- | --- | --- |
| | Biomass<br>$X_{cin}$ | Biomass<br>$X_{thr}$ | Aqueous<br>$S_{Ac}$ | |
| 1. *Methanosarcina* Growth | +1 | 0 | $-1/Y_{cin}$ | $\mu_{m,cin} \frac{S_{Ac}}{S_{Ac}+K_{Ac,cin}} X_{cin}$ |
| 2. *Methanosarcina* Decay | -1 | 0 | 0 | $b_{cin} X_{cin}$ |
| 3. *Methanosarcina* Outflow | -1 | 0 | 0 | $\frac{X_{cin}}{\theta_x}$ |
| 4. *Methanothrix* Growth | 0 | +1 | $-1/Y_{thr}$ | $\mu_{m,thr} \frac{S_{Ac}}{S_{Ac}+K_{Ac,thr}} X_{thr}$ |
| 5. *Methanothrix* Decay | 0 | -1 | 0 | $b_{thr} X_{thr}$ |
| 6. *Methanothrix* Outflow | 0 | -1 | 0 | $\frac{X_{thr}}{\theta_x}$ |
| 7. $S_{Ac}$ Inflow | 0 | 0 | +1 | $\frac{S_{Ac}^0}{\theta_x}$ |
| 8. $S_{Ac}$ Outflow | 0 | 0 | -1 | $\frac{S_{Ac}}{\theta_x}$ |

**Extended Data Table 2:** Microbial kinetic parameters for *Methanosarcina* and *Methanothrix* for acetoclastic methanogenesis at mesophilic conditions based on previous compilations. This work uses the midpoint of the range.

| Parameter (unit) | *Methanosarcina* | *Methanothrix* | References |
| --- | --- | --- | --- |
| *Literature range* |  |  |  |
| $\mu_m$ (d<sup>-1</sup>) | 0.20 – 1.4 | 0.09 – 0.72 | 17 |
| $K_{Ac}$ (meeq<sub>ac</sub> L<sup>-1</sup>)<sup>a</sup> | 18 – 46 | 1.3 – 11 | 16,17 |
| $Y$ (meeq<sub>X</sub> meeq<sub>ac</sub><sup>-1</sup>) | 0.026 – 0.14 | 0.023 – 0.045 | 17 |
| $b$ (d<sup>-1</sup>) | 0.004 – 0.10 | 0.006 – 0.020 | 17 |
| *Parameters selected for this work (midpoint)* |  |  |  |
| $\mu_m$ (d<sup>-1</sup>) | 0.80 | 0.41 | |
| $K_{Ac}$ (meeq<sub>ac</sub> L<sup>-1</sup>) | | | |
| Regimes I to IV | 32 | 6.3 |  |
| Regimes V to VI | 32 | 22<sup>b</sup> |  |
| $Y$ (meeq<sub>X</sub> meeq<sub>ac</sub><sup>-1</sup>) | 0.083 | 0.034 | |
| $b$ (d<sup>-1</sup>) | 0.052 | 0.013 | |

<sup>a</sup> Converted from mg COD L<sup>-1</sup> using 8 mg COD = 1 meeq (COD stands for chemical oxygen demand).
<sup>b</sup> Adjusted to include thermodynamic inhibition and be consistent with $S_{min}^*$ estimated by McCarty and Bae<sup>33</sup>.

**Extended Data Table 3:** Reaction stoichiometry and the standard Gibbs free energy for anabolic, catabolic, and endogenous decay reactions for acetoclastic methanogenesis. All reactions are written in terms of 1 electron equivalent (eeq) transferred. Reactions are defined at pH=7, T=298.15K, and unit activity.

| **Reaction Name** | **Stoichiometry** | **$\Delta G^{o'}$**<br>**[kJ/eeq]** |
| --- | --- | --- |
| Anabolic | $0.125 \text{ CH}_3\text{COO}^- + 0.05 \text{ NH}_4^+ + 0.075 \text{ H}^+ \rightarrow 0.05 \text{ C}_5\text{H}_7\text{O}_2\text{N} + 0.15 \text{ H}_2\text{O}$ | 6.96 |
| Catabolic | $0.125 \text{ CH}_3\text{COO}^- + 0.125 \text{ H}^+ \rightarrow 0.125 \text{ CH}_4 + 0.125 \text{ CO}_2$ | -4.48 |
| Endogenous Decay | $0.05 \text{ C}_5\text{H}_7\text{O}_2\text{N} + 0.15 \text{ H}_2\text{O} + 0.05 \text{ H}^+ \rightarrow 0.125 \text{ CH}_4 + 0.05 \text{ NH}_4^+ + 0.125 \text{ CO}_2$ | -11.44 |

The standard Gibbs energy of formation is taken from Heijnen and Kleerebezem<sup>34</sup> for all compounds, except for biomass. Biomass is represented by $\text{C}_5\text{H}_7\text{O}_2\text{N}$ with the Gibbs free energy of formation $\Delta G_f^{o'} = 35 \text{ kJ eeq}^{-1}$ , which falls near the midrange of compiled experimentally measured values Torres et al.<sup>35</sup>. $\Delta G_{ana}^{o'}$ value is consistent with Chang et al.<sup>17</sup>.

**Extended Data Table 4:** Energy dissipation matrix for the two-species acetoclastic methanogenesis chemostat model. *Methanosarcina* (cin) and *Methanothrix* (thr) abbreviations follow Extended Data Table 1. The Gibbs free energy of reaction, Δ*G* (kJ per eeq_*rxn*_), is for reactions defined in Extended Data Table 3 after concentration corrections.

| Process | $\Sigma$ | | $\Delta G$ | | $\rho(\mathbf{E})$ |
| --- | --- | --- | --- | --- | --- |
| | $\sigma_{p,ana}$ | $\sigma_{p,cat}$ | $\Delta G_{p,ana}$ | $\Delta G_{p,cat}$ | |
| 1. *Methanosarcina* Growth | 1 | $(1 - Y_{cin})/Y_{cin}$ | $\Delta G_{grow,ana,cin}$ | $\Delta G_{grow,cat,cin}$ | $\mu_{m,cin} \frac{S_{Ac}}{S_{Ac} + K_{Ac,cin}} X_{cin}$ |
| 2. *Methanosarcina* Decay | 0 | 1 | 0 | $\Delta G_{dec,cat}$ | $b_{cin} X_{cin}$ |
| 3. *Methanothrix* Growth | 1 | $(1 - Y_{thr})/Y_{thr}$ | $\Delta G_{grow,ana,thr}$ | $\Delta G_{grow,cat,thr}$ | $\mu_{m,thr} \frac{S_{Ac}}{S_{Ac} + K_{Ac,thr}} X_{thr}$ |
| 4. *Methanothrix* Decay | 0 | 1 | 0 | $\Delta G_{dec,cat}$ | $b_{thr} X_{thr}$ |

**Extended Data Figure 1.**
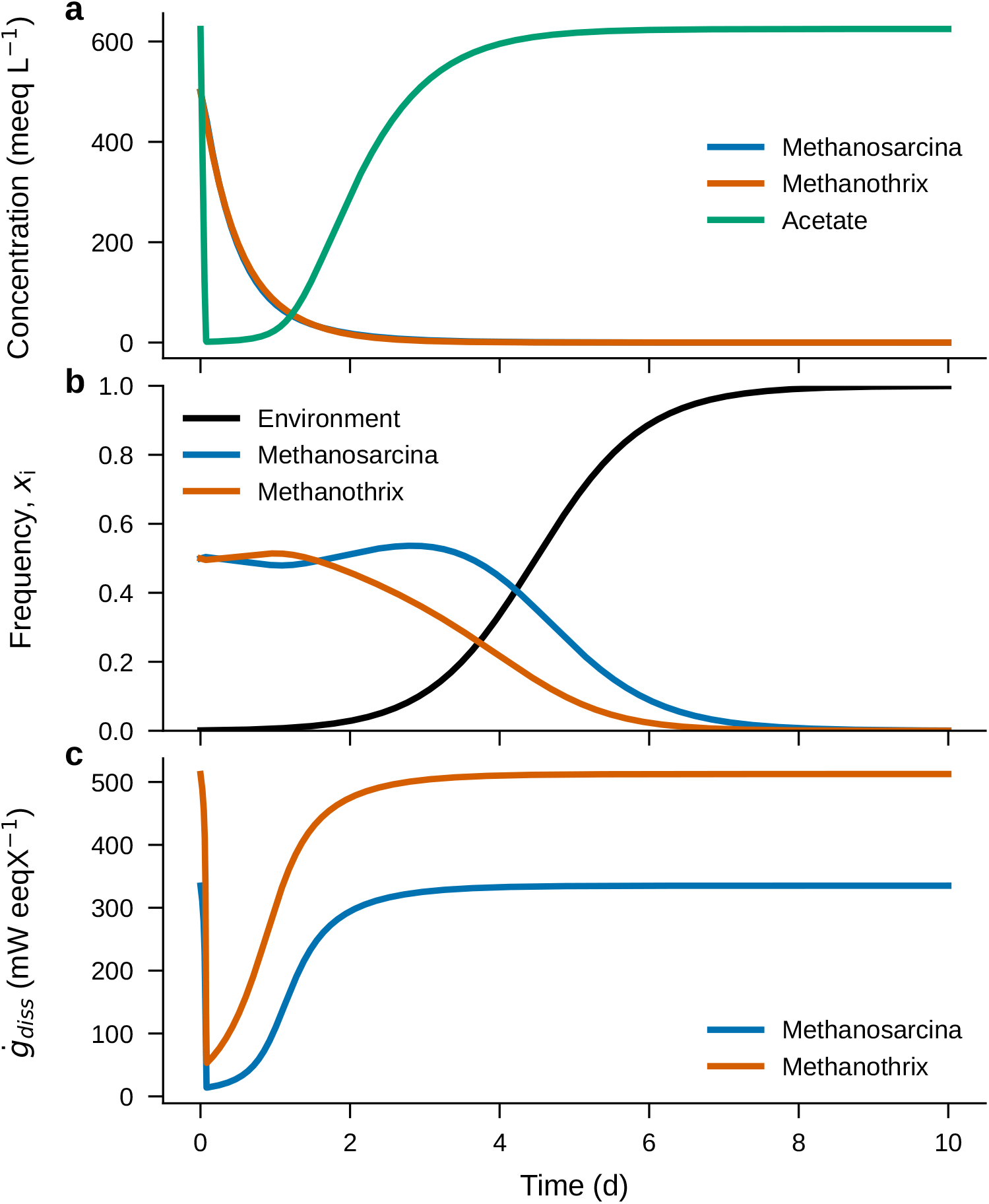
Illustration of Washout (Regime I) in (a) absolute coordinates, (b) the simplex, and (c) specific energy dissipation rate. Parameters are summarized in Extended Data Table 2. Initial conditions are *X*_*cin*_ = 500 meeq L−1, *X*_*thr*_ = 500 meeq L−1, and *S*_*Ac*_ = 625 meeq L−1.

**Extended Data Figure 2.**
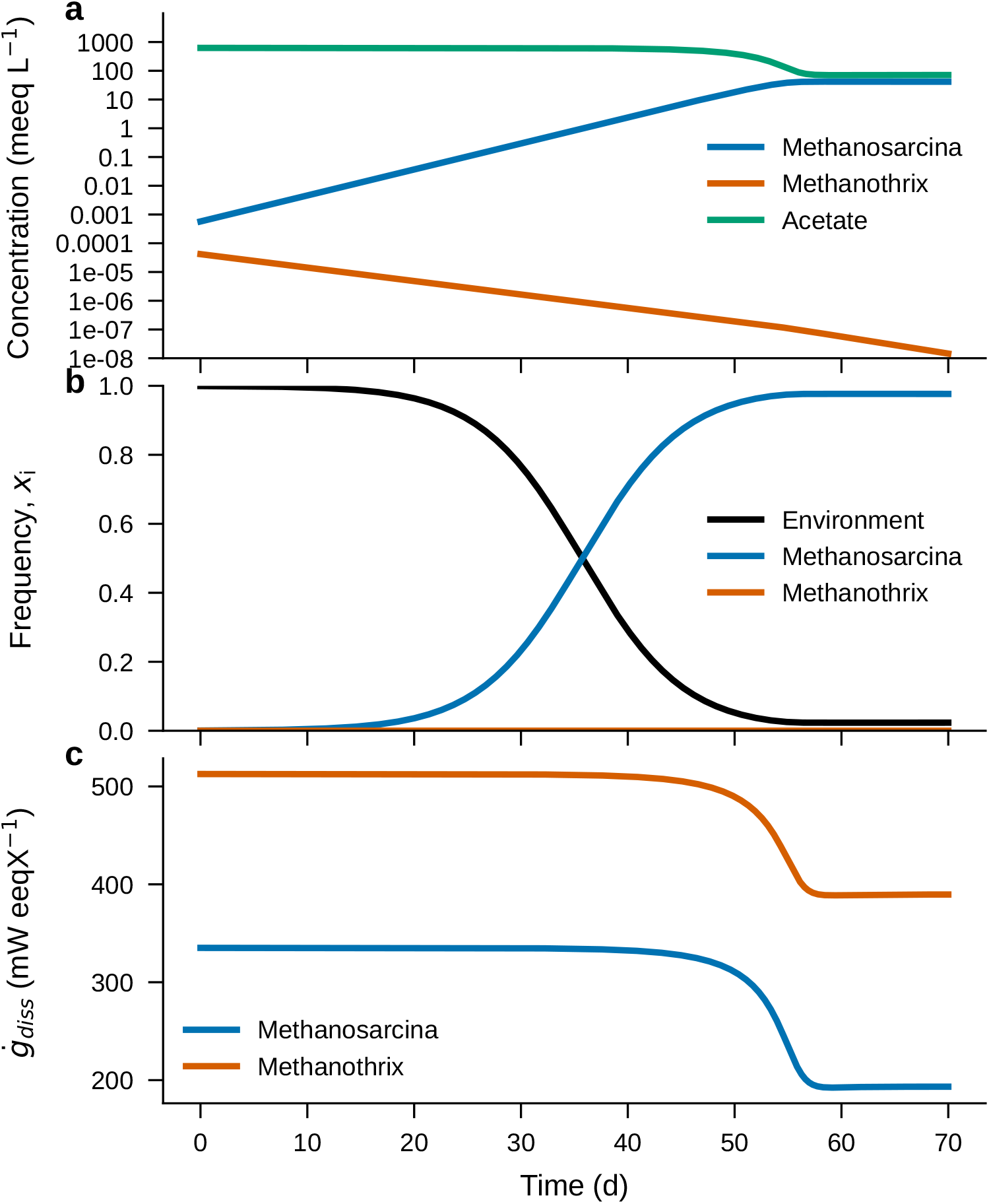
Illustration of r-dominates (Regime II) in (a) absolute coordinates, (b) the simplex, and (c) specific energy dissipation rate. Parameters are summarized in Extended Data Table 2. Initial conditions are *X*_*cin*_ = 5.7 × 10−4 meeq L−1, *X*_*thr*_ = 4.2 × 10−5 meeq L−1, and *S*_*Ac*_=625 meeq L−1. The y-axis is shown in the log scale for Panel a)

**Extended Data Figure 3.**
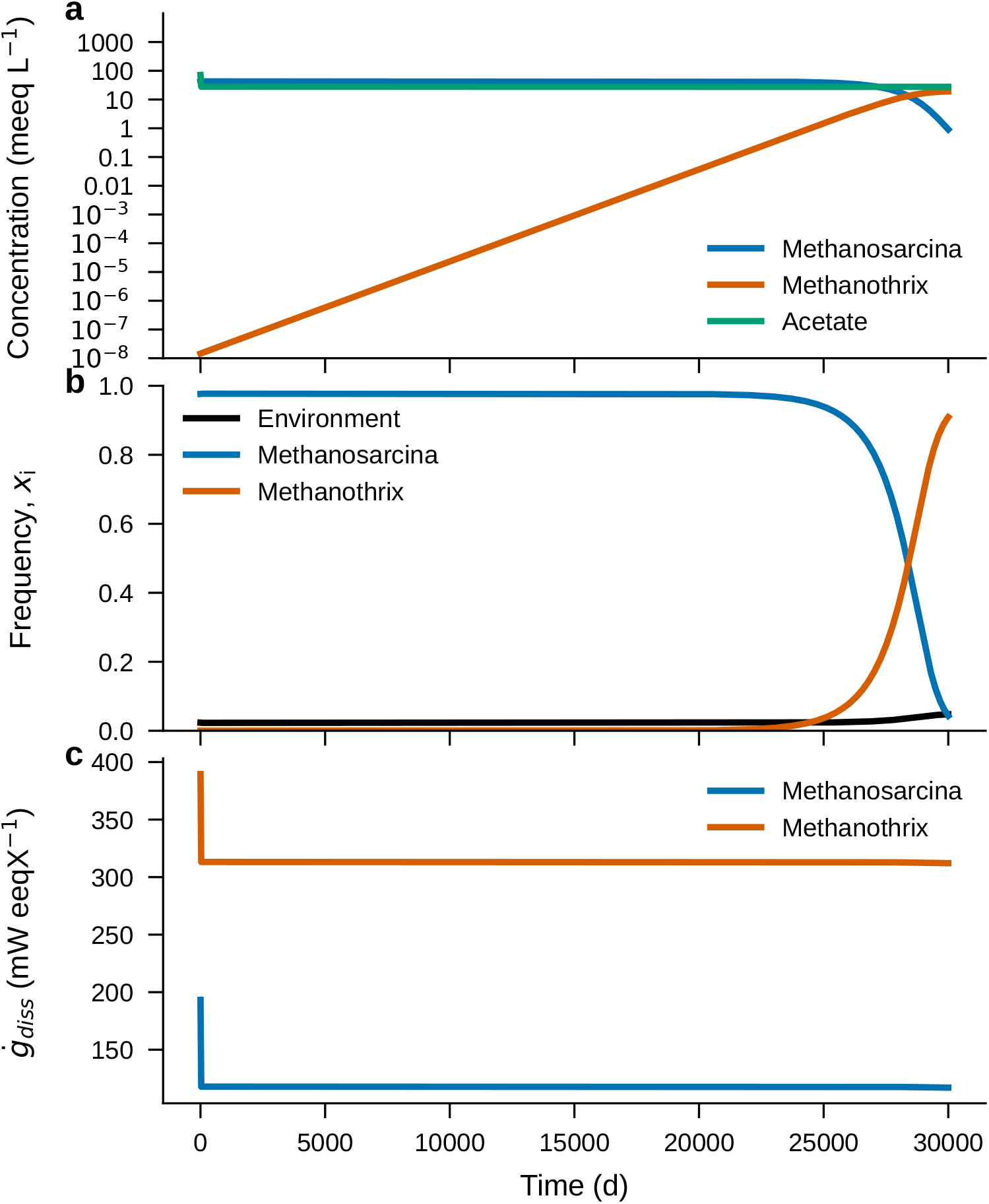
Illustration of K-dominates (Regime IV) in (a) absolute coordinates, (b) the simplex, and (c) specific energy dissipation rate. Parameters are summarized in Extended Data Table 2. Initial conditions are *X*_*cin*_ = 42 meeq L−1, *X*_*thr*_ = 1.4 × 10−8 meeq L−1, and *S*_*Ac*_=71 meeq L−1. The y-axis is shown in the log scale for Panel a).

**Extended Data Figure 4.**
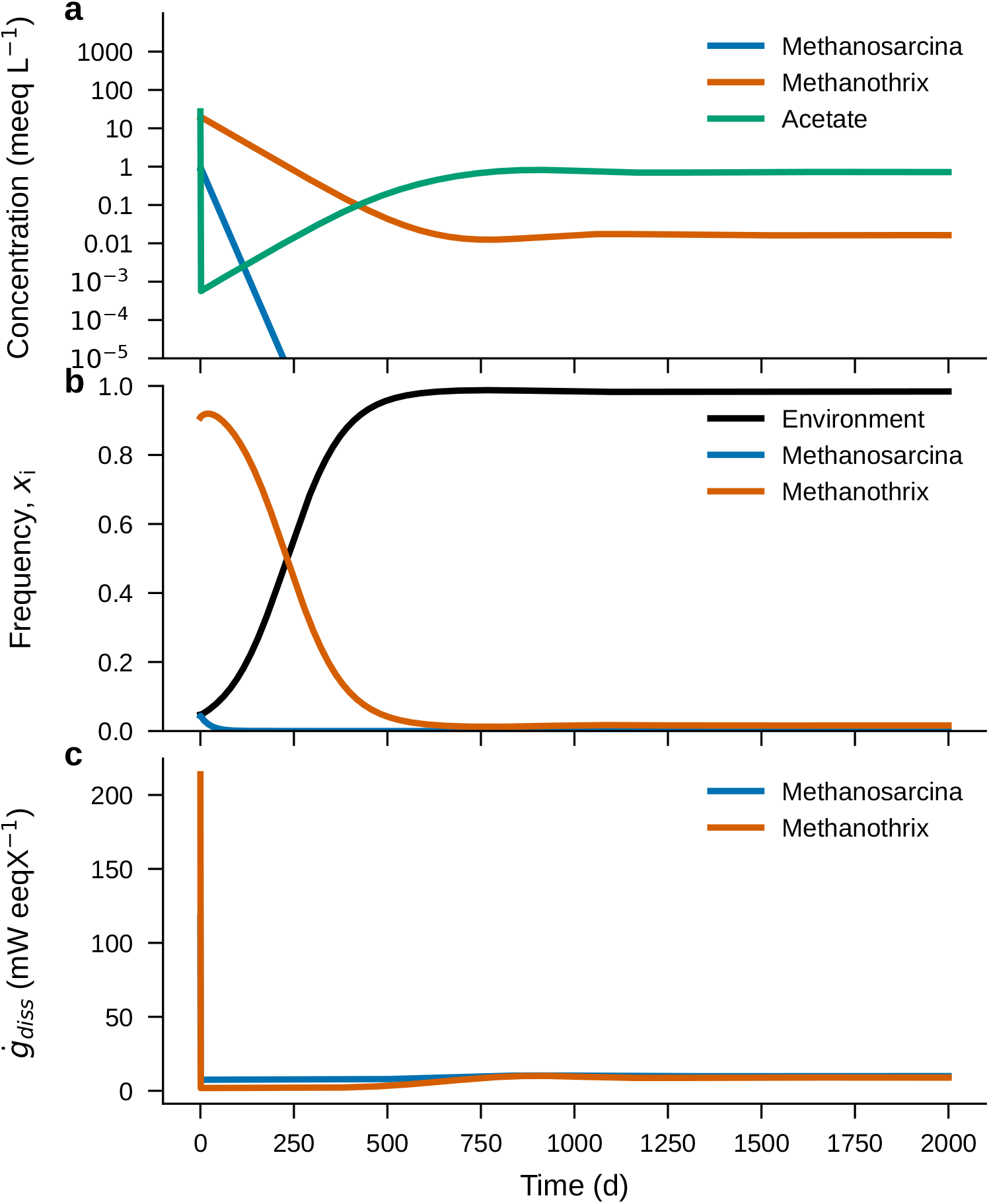
Illustration of 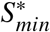 (Regime V) in (a) absolute coordinates, (b) the simplex, and (c) specific energy dissipation rate. Parameters are summarized in Extended Data Table 2. Initial conditions are *X*_*cin*_ = 0.034 meeq L−1, *X*_*thr*_ = 20 meeq L−1, and *S*_*Ac*_=28 meeq L−1. The y-axis is shown in the log scale for Panel a).

## S1 The Grand Vision

Few topics inspire more Greatest of All Time (GOAT) debates than the rivalry among Roger Federer, Rafael Nadal, and Novak Djokovic. Federer represents a strategy of an elegant, but truly aggressive, all-court offense. Nadal represents a strategy of physical, heavy-spin baseline play. Djokovic represents a strategy of the ultimate counterpunching. Over the past two decades, these three legends have combined 66 grand slam wins. Some may argue that their competition has reached an apparent stalemate – no single playing style has a decisive advantage over the others. Who wins on any given day may depend on chance alone.

Is there a way to break the stalemate? An avid sports fan may argue that the court environment can break the stalemate: the fast grass of Wimbledon favors Federer’s strategy, the slow clay court of Roland Garros favors Nadal’s strategy, and the medium-speed hard courts of the Australian Open and the US Open favor Djokovic’s strategy. If the environment is considered, should Pete Sampras be in the conversation? Sampras represents a classic serve-and-volley strategy and dominated in the era before Wimbledon changed its grass courts. How about John McEnroe and Björn Borg, who played with wooden rackets? These examples suggest that a proper evaluation of each strategy requires an explicit consideration of the evolving environments.

This Supplementary Note develops a mathematical foundation that can handle the GOAT debate by introducing the environment as a player in a game of strategy.

The Supplementary Note is written with the recognition that the Environmental Games rest on two pillars, each coming from two separate traditions. Evolutionary Game Theory (EGT) has its historical origins in classical game theory by von Neumann and Morgenstern^33^ and Nash^34^, where players make rational decisions and mix strategies to optimize their outcomes – like how Federer may change the ratio of lob shots and passing shots depending on who he plays. Independent of the game theory tradition, Smith and Price^9^ developed the concept of evolutionarily stable strategy (ESS) to describe whether a majority in a population can adopt a strategy to resist invasion from a rare mutant. The replicator equation was developed by Taylor and Jonker^35^ to connect the static equilibrium concepts of Price and Smith to a dynamical process. In replicator dynamics, strategies are hardwired into individual phenotypes, unlike rational agents making decisions. This framework is particularly well-suited for modeling microbiological systems, where microbes have fixed phenotypes instead of acting like rational agents. Comprehensive treatments of EGT are given by Hofbauer and Sigmund^1^, Nowak^3^.

Environmental Biotechnology is an engineering discipline that applies principles from reactor engineering and microbial ecology to grow microorganisms that benefit society^11,13^. Environmental Biotechnology is a forward-looking science that sets the right environmental conditions and lets the beneficial microorganisms already present in the environment grow into abundance. Perry McCarty laid many of the early foundations of thermodynamics and reactor engineering principles governing environmental selection^36,37^.

Readers will likely be familiar with at most only one of the two above-mentioned pillars. Therefore, the Supplementary Note provides a detailed derivation that engineers and microbiologists may find useful. A caveat is that the tennis example above is meant to illustrate the game of interactions within the framework of classical game theory, where each player makes rational decisions. The Environmental Games will build on the framework of replicator dynamics, where each strategy is represented by distinct population frequencies.

The remaining sections introduce the replicator equation and EGT terminology (S2). Then, the equivalence between the replicator equation and the Lotka-Volterra model (LVM) (S3) is demonstrated following Hofbauer and Sigmund^1^ . This mathematical procedure is essential for projecting the absolute coordinates of the LVM and Environmental Biotechnology models onto the simplex described by the replicator equation. It also separates direct interaction coefficients and intrinsic growth rates within the payoff matrix. Setting the direct interaction coefficients to zero yields the idealized LVM (S4) – a baseline selection model with no direct biological interactions. The Pure Environmental Games (PEGs) are derived by replacing the intrinsic growth rate with the net specific accumulation rate of biomass (S5). Finally, the classical r/K selection game will be used as an instance of PEGs, and six environmental selection regimes will be described (S6).

## S2 Replicator Dynamics

Classical game theory provides a formal framework for studying strategic interactions among players trying to maximize their rewards. In a game with two strategies, a player can select from one of two “pure” strategies *S*_1_ and *S*_2_. The payoff coefficient *a*_*i j*_ describes the payoff for strategy *i* interacting with *j*. One way to show the payoff matrix is to let the rows indexed by *i* represent the focal strategy and the columns indexed by *j* represent the opposing strategy. A payoff matrix for a general game with two strategies, with the coefficients *a*_11_ = *a, a*_12_ = *b, a*_21_ = *c, a*_22_ = *d*, can be shown as

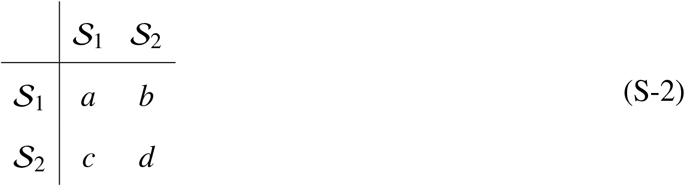

where a, b, c, and d are defined on R. In practice, a player will play a mixture of *S*_1_ and *S*_2_ and this mixed strategy can be described using a probability distribution: *P*(*S*_1_) and *P*(*S*_2_). By the law of total probability *P*(*S*_1_) + *P*(*S*_2_) = 1. Let us assume that the opponent plays *S*_1_ and *S*_2_ with probabilities *P*(*S*_1_) and *P*(*S*_2_). Then, the net payoff for the player playing *S*_1_ is

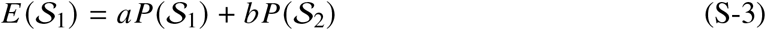

and *S*_2_ is

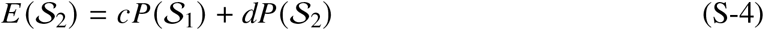

These are naive calculations assuming an arbitrary probability distribution before optimization. In classical game theory, the optimal strategies are developed by anticipating the opponent’s optimal moves, which depend on how the games are played (finite versus infinite rounds). For a complete treatment of classical game theory, readers are referred to Nowak^3^, von Neumann and Morgenstern^33^.

The replicator equation is a fundamental deterministic framework for modeling large, wellmixed populations in which individuals undergo repeated interactions. In contrast to classical game theory, where rational players choose strategies, each individual represents a specific, fixed strategy (phenotype). If a population of size N has *k*_*i*_ individuals playing a strategy *i*, the frequency of strategy *i* is defined as

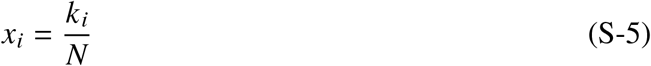

The population frequencies for a population with n+1 strategies sum to one: 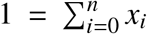. In replicator dynamics, each strategy undergoes continuous frequency-dependent interactions to shape the population.

In replicator dynamics, the benefits or losses that each individual receives from another individual in a population are calculated in a manner that is analogous to classical game theory, but with important differences: The strategy frequency *x*_*i*_ replaces probability, and the fitness *f*_*i*_ replaces the net payoff. For a two-strategy case, the fitness of two strategies can be calculated as

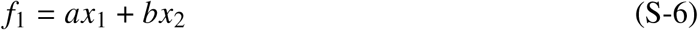

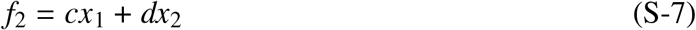

or more generally

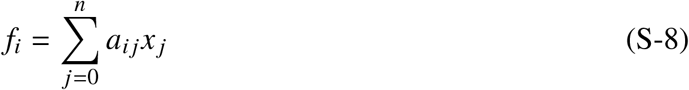

For non-mathematicians unfamiliar with the summation notation, notice the index *j* is summed over, leaving *i* as the matching index for both *a*_*i j*_ and *f*_*i*_. While the mathematical forms between the net payoff and fitness are analogous, the interpretation differs fundamentally. In classical game theory, mixing reflects a single player’s probabilistic choice, whereas in replicator dynamics, mixing reflects the population composition of individuals each committed to a fixed pure strategy.

The replicator equation is convenient for studying large mixed populations, because it has a built-in mechanism to track the outcome of repeated interactions. It models the frequency of strategy *i* in a population (*x*_*i*_) over continuous and repeated interactions in the simplex space and has the form:

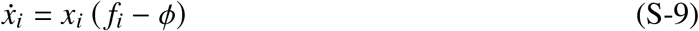

where *ϕ* is the average fitness of the population, and

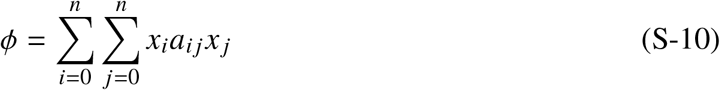

Note: this work adopts a zero-indexing convention, with *x*_0_ denoting the empty strategy, as discussed later. The replicator equation is a remarkably intuitive equation that says the frequency of a strategy will increase if its own fitness exceeds that of the population average.

The payoff matrix can be interpreted through two different classification systems: The ESS classification and the EGT classification. The ESS classification describes static equilibria and asks if a strategy can resist an invasion by a rare mutant by examining the payoff matrix3. Consider a game of direct interactions with two strategies: *S*_1_ and *S*_2_, where a payoff matrix for a 2-strategy system with *a*_11_ = 1, *a*_12_ = 0, *a*_21_ = 0, *a*_22_ = 0 can be shown as

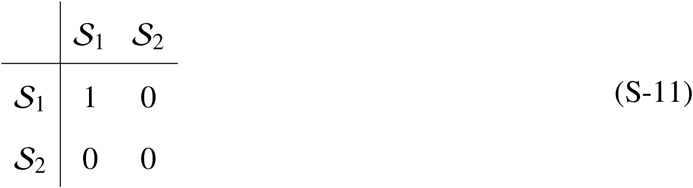

A strategy is a strict Nash equilibrium (SNE) if it resists all invasions. The strategy *S*_*k*_ is an SNE if it satisfies

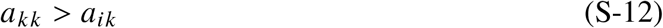

for all *i*. In this example, *S*_1_ is an SNE because the diagonal element in the first column is larger than the other elements in the same column: i.e., *a*_11_ > *a*_21_ or 1 > 0. An ESS also resists all invasions from emerging mutants. An SNE implies ESS, but ESS for *S*_*k*_ can be established by a less stringent criterion:

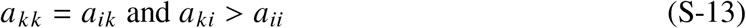

for all *i*. Although *S*_1_ fails the second criterion (i.e., *a*_12_ ≯ *a*_22_ or 0 ≯ 0), it is saved by SNE already being satisfied: *S*_1_ is both SNE and ESS.

Let’s see if *S*_2_ satisfies ESS. Because *a*_22_ = *a*_12_ or 0 = 0, the first part of ESS criteria is satisfied. However, since *a*_21_ ≯ *a*_11_ or 0 ≯ 1, *S*_2_ is not an ESS. So, what is *S*_2_? The answer is in the Nash equilibrium (NE). The criteria for the NE are

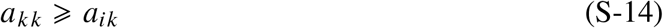

Because *a*_22_ = *a*_12_ or 0 = 0, *S*_2_ is an NE. An NE does not prevent invasion, but it offers a possibility for neutral drift.

SNE, ESS, and NE are static equilibrium concepts describing the ability of a resident strategy to resist invasion from a rare strategy. These concepts need to be distinguished from the following concepts:

- In EGT, dynamic equilibrium describes a state when the vector of frequencies becomes time invariant. This state can be stable or unstable.
- A related concept in Environmental Biotechnology is the steady state, which occurs when the environmental variables, such as concentrations of substrate and biomass, become time invariant.
- A thermodynamic equilibrium which occurs when the overall Gibbs free energy of reactions equals zero (Δ*G*_*rxn*_ = 0).

The EGT classification or the game given by the payoff structure describes the categories of interactions demonstrated by strategies. Table S1 below shows the correspondence between the game and replicator dynamics for a two-strategy game. Traditionally, replicator dynamics assume that the intrinsic growth rates of individual strategies are equal. The game and replicator dynamics have a one-to-one correspondence when the intrinsic growth rates are equal^2^. The PEGs explore differences in intrinsic growth rates operating in parallel to the limiting case of neutral selection, revealing that deterministic outcomes remain possible even when direct interactions are absent. The dynamics of frequency-based selection are discussed in Supplementary Note S4.

**Supplementary Table S1:** EGT classification of two-strategy games, corresponding replicator dynamics, and payoff structure conditions. The one-to-one correspondence between game and replicator dynamics holds when intrinsic growth rates are equal^2^.

| Game | Replicator Dynamics | Payoff Structure |
| --- | --- | --- |
| Harmony Game | Dominance of $\mathcal{S}_1$ | $a > c$ and $b > d$ |
| Prisoner's Dilemma | Dominance of $\mathcal{S}_2$ | $a < c$ and $b < d$ |
| Stag Hunt | Bistability | $a > c$ and $b < d$ |
| Hawk-and-Dove | Coexistence | $a < c$ and $b > d$ |
| Neutral Selection | Neutral | $a = c$ and $b = d$ |

## S3 Transforming a Two-species Lotka-Volterra System into a Replicator Equation

### The Lotka-Volterra Systems

The LVM from theoretical ecology also describes the game of direct interactions. Instead of the relative frequency, the LVM tracks the absolute abundance of individual *i* in a population (*y*_*i*_). The LVM has the form:

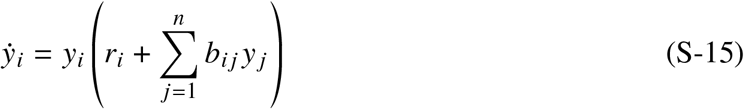

*r*_*i*_ stands for intrinsic growth rate of species *i*, and *b*_*i j*_ stands for gains or losses species *i* experiences from interacting with species *j*. Note that the LVM adopts a one-indexing convention. Hofbauer and Sigmund^1^ demonstrate that the LVM with n species is mathematically equivalent to the replicator equation with n + 1 strategies. However, there are important physical differences between these two frameworks. On the one hand, the replicator equation only has one set of coefficients in the payoff matrix that represents direct player interactions. On the other hand, the LVM has two sets of coefficients: the direct interaction coefficients and the intrinsic growth rates. This section transforms a two-species Lotka-Volterra system to the replicator equation to demonstrate how the coefficients of the Lotka-Volterra system map onto the payoff matrix.

Consider a two-species Lotka-Volterra system in the form:

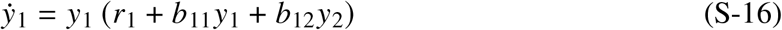

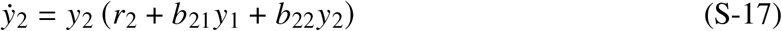

The interaction terms and intrinsic growth rates for the two-species Lotka-Volterra system can be represented in the matrix form as

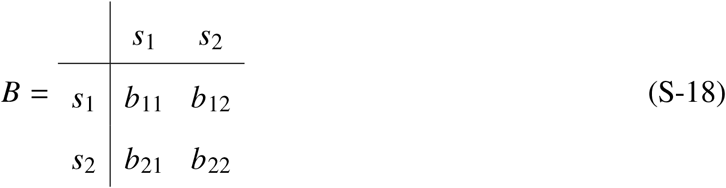

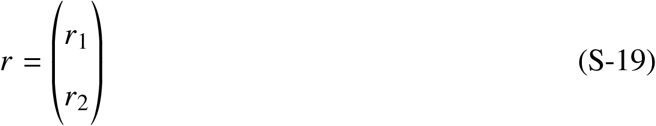

where *s*_1_ and *s*_2_ represent species. The species abundance is defined on R^+^.

### Folding Infinity onto a Simplex

Transforming the LVM into the replicator equation requires a map between absolute quantities defined over [0, ∞) and frequencies bounded within a simplex: i.e., a set of points whose coordinates are non-negative and sum to one^3^. An intuitive approach is using the total population density as the normalization factor. In Environmental Biotechnology and biofilm modeling, this mapping is routinely achieved by normalizing the individual biomass density by the total biomass density to define the biofilm fraction^38,39^. This is not a desirable transformation approach because the replicator equation does not track absolute quantities.

An effective alternative is to introduce a normalization factor embedded with a unit reference value. Hofbauer and Sigmund^1^ define the normalization factor *D* as

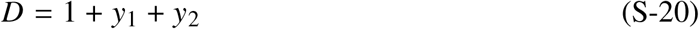

The value 1 in (S-20) represents a unit value carrying the same dimension as *y*_*i*_, analogous to a reference concentration of 1 mol/L in chemical systems. This method maps the absolute quantities to a coordinate in the simplex frequencies, *x*_*i*_, as

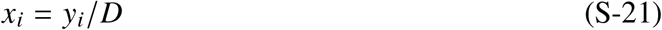

Because *D* carries the same physical dimension as *y*_*i*_, *x*_*i*_ is dimensionless. Using this mapping, (S-20) can be divided by *D* and rewritten as

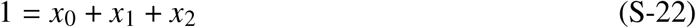

It should be noted that *x*_0_ = 1/*D* and substituting this relationship to (S-20) gives

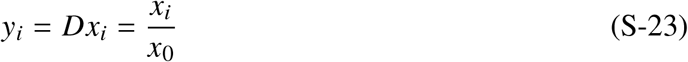

*x*_0_ is defined over (0, 1]. When the simplex is unoccupied (Σ*i y*_*i*_ = 0), *x*_0_ = 1. *x*_0_ diminishes as pure strategies occupy the simplex. When the absolute quantity approaches infinity (Σ*i y*_*i*_ →− ∞), *x*_0_ approaches zero. Thus, this mapping preserves the relationship between simplex coordinates and absolute coordinates by dynamically tracking *x*_0_.

As noted earlier, this method increases the number of state variables from n in the LotkaVolterra system to n+1 in replicator dynamics. Since *x*_0_ does not represent any species in the original Lotka-Volterra system, it can be thought of provisionally as the frequency of an empty strategy, *S*_0_. However, Supplementary Note S5 shows that *x*_0_ represents the environment as the player in the Pure Environmental Games.

### The Transformation

With the empty strategy, the two-species Lotka-Volterra system can now be expressed as replicator equations. The transformation proceeds by taking logarithmic derivatives. Taking the logarithmic derivatives of (S-21)

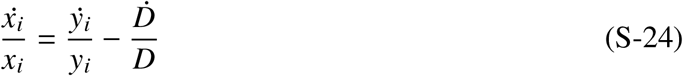

Since *y*_0_ = 1, (S-21) for the empty strategy is

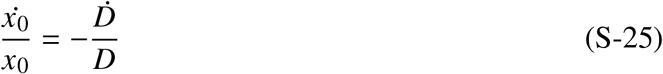

The “dot” notation in 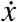 represents the derivative with respect to dt. By defining a new coordinate system, 1/*dτ* = *x*_0_/*dt*, the average fitness *ϕ* is defined as

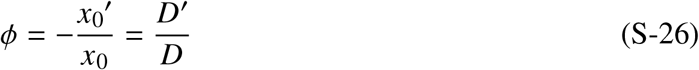

This expression can be rearranged to the replicator equation for the empty strategy:

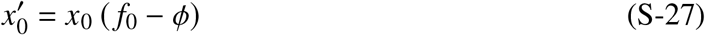

where it is useful to note that the fitness of the empty strategy is *f*_0_ = 0. Moreover, inserting (S-16) and (S-25) into (S-24) gives

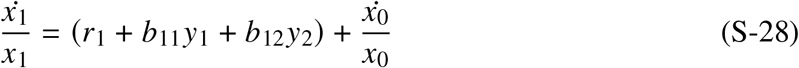

The terms *y*_1_ and *y*_2_ in this equation can be eliminated by noting *y*_*i*_ = *x*_*i*_/*x*_0_ from (S-21)

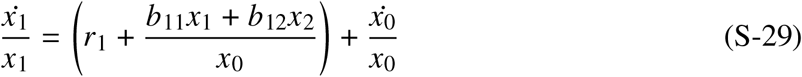

Multiplying the equation by *x*_0_ gives

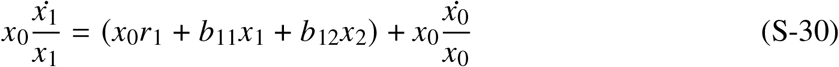

Employing the same time domain transformation, 1/*dτ* = *x*_0_/*dt*, gives

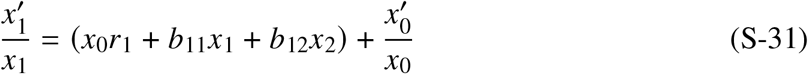

where prime indicates differentiation with respect to the transformed time coordinate, 1/*dτ* = *x*_0_/*dt*. Noticing that the terms in the parenthesis in the fitness of species 1 and the definition of average fitness in (S-25) provide the replicator equation for species 1

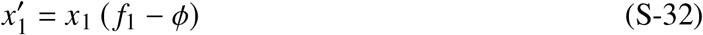

where

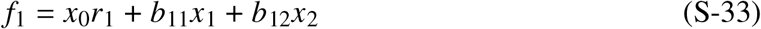

Similarly, the replicator equation for species 2

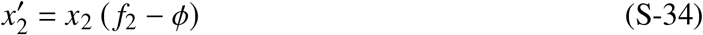

where

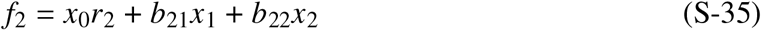

Together (S-27), (S-32), and (S-34) provide the three replicator equations from the two-species Lotka-Volterra system. The average fitness can be obtained by summing these three replicator equations to obtain

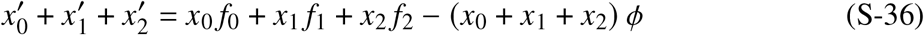

Notice from (S-22) that *x*_0_ + *x*_1_ + *x*_2_ = 1 and its derivative is zero: i.e., 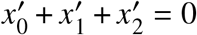. Therefore, obtain the average fitness

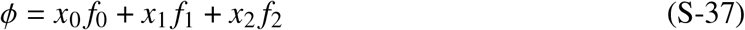

This concludes the transformation of the two-species Lotka-Volterra system into equivalent replicator equations. The resulting coefficient for the replicator equation can be summarized in a payoff matrix:

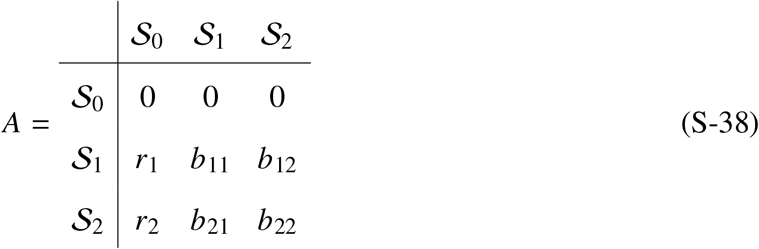

The payoff matrix derived from the Lotka-Volterra system has several important features. First, now that we are in the simplex space, *S*_*i*_ stands for strategies, not species. *S*_1_ and *S*_2_ are pure strategies adopted by *s*_1_ and *s*_2_. Meanwhile, *S*_0_ is an empty strategy introduced to act as the reference size of the simplex (S-20). Second, the first row contains zero because the empty strategy receives no payoff. Third, the second and third columns contain the interaction terms from the two-species Lotka-Volterra systems. Fourth, the first column contains the intrinsic growth rates from the Lotka-Volterra system. Conceptually, each term represents the payoff a strategy receives by playing against an empty strategy. When the payoff matrix operates on the strategy frequencies, the term *x*_0_ multiplies the intrinsic terms. Therefore, *x*_0_ acts as a conversion factor to convert the intrinsic growth rates of Lotka-Volterra system into an equivalent fitness value in replicator dynamics. The next section shows how *x*_0_ provides the link to incorporate the boundary condition, such as the carrying capacity, in the simplex space when considering a system derived from LVM. The subsequent sections demonstrate how *x*_0_ provides the link to bring physical and chemical constraints into the simplex space.

## S4 The Ideal Lotka-Volterra Systems

### Predator-Prey System

Theoretical ecologists use the LVM to study interactions. A classic example is the Lotka-Volterra Predator-Prey model of the form

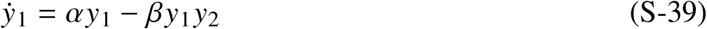

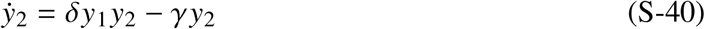

where *α* is prey birth rate, *γ* is predator death rate, and *β* and *δ* are interaction coefficients. Using the results from transforming the two-species Lotka-Volterra systems to the replicator equation in Supplementary Note S3 (S-38), the coefficients for the Predator-Prey system can be placed in the payoff matrix as shown below:

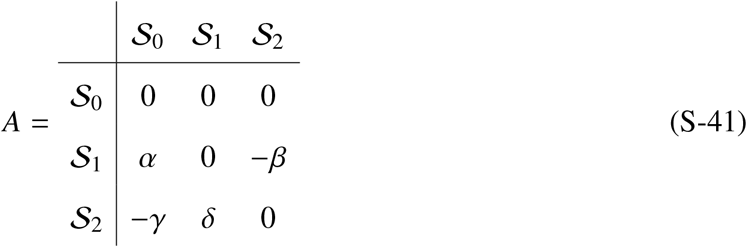

There are a few notable features. First, the first column representing the empty strategy contains the intrinsic growth terms. As stated in Supplementary Note S3, *x*_0_ re-denominates *α* and *γ* from absolute units into simplex frequencies to alter the fitness of *S*_1_ and *S*_2_, respectively. Then, because the payoff coefficients are defined on R, the term *γ* carries the negative sign from the equation. Next, the interaction terms *β* and *δ* encode the predator-prey interactions. The term *β* carries a negative sign because predation results in the prey’s loss. Finally, zero diagonal entries indicate no self-interaction.

Supplementary Figure S1a shows the model dynamics when coefficients equal *α* = 1.0, *β* = 0.1, *δ* = 0.075, *γ* = 1.5. This system has a non-trivial equilibrium at 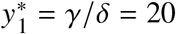 and 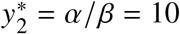. When the system starts at an initial point away from this equilibrium (*y*_1_ = 25, *y*_2_ = 10 in this example), the population dynamics demonstrate an oscillatory behavior around the equilibrium.

Supplementary Figure S1b shows the model dynamics in the simplex. In the simplex, the x-axis is represented by the transformed time coordinate, *τ*. The simplex adds the empty strategy to track changes in absolute abundance through *x*_0_.

### Ideal Lotka-Volterra

Let us define the Ideal LVM in which no individuals within the population interact directly (similar to the ideal gas law in chemistry). The model is also known as the Malthusian growth model or simply the exponential growth model. We shall see in the coming sections that it becomes convenient to strip away the interaction terms while preserving the structure of the LVM. Mathematically, the LVM is represented by setting all interaction coefficients in (S-15) to zero. Consider an ideal two-species Lotka-Volterra system in the form:

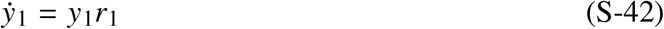

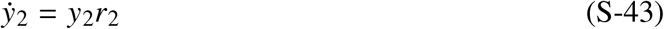

When this ideal Lotka-Volterra system is projected onto the simplex, the following is the payoff matrix:

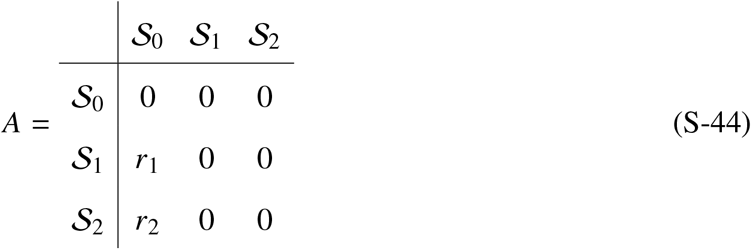

The ESS classification can be applied to the ideal Lotka-Volterra system to identify when the empty strategy can be invaded.

- When *r*_1_, *r*_2_ > 0, none of the strategies is an SNE because *a*_*kk*_ ≯ *a*_*ik*_ (S-12).
- When *r*_1_, *r*_2_ > 0, none of the strategies is an ESS because although *S*_1_ and *S*_2_ satisfy *a*_*kk*_ = *a*_*ik*_, *a*_*ki*_ ≯ *a*_*ii*_ (S-13).
- *S*_1_ and *S*_2_ are always Nash equilibrium strategies because *a*_*kk*_ ⩾ *a*_*ik*_ or 0 ⩾ 0 (S-14), regardless of the signs of *r*_1_ and *r*_2_.
- When *r*_1_, *r*_2_ > 0, *S*_0_ does not qualify for a Nash equilibrium (S-14) because *a*_00_ = 0 ≯ *a*_10_ = *r*_1_ > 0.
- Interestingly, when *r*_1_, *r*_2_ < 0, the empty strategy becomes an SNE because *a*_00_ > *a*_10_ and *a*_20_ (S-12).

**Supplementary Figure S1:**
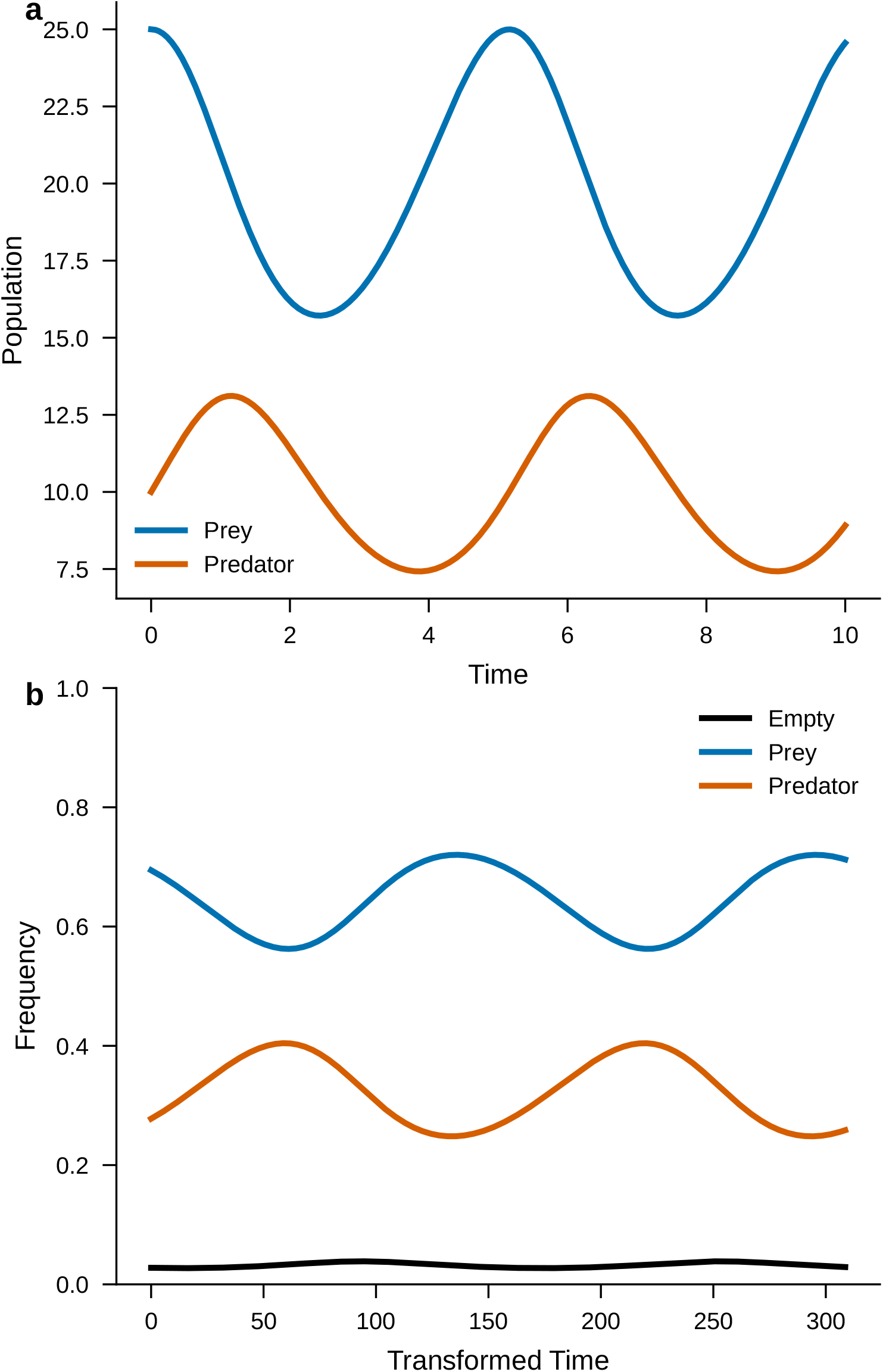
Predator-prey dynamics in a) absolute coordinates and b) simplex coordinates. Parameters: *α* = 1.0, *β* = 0.1, *δ* = 0.075, *γ* = 1.5. Initial conditions: prey = 25, predator = 10.

Therefore, the ideal LVM consists of a collection of Nash equilibrium strategies that can successfully establish themselves in the simplex by displacing the empty strategy when *r*_1_, *r*_2_ > 0. However, when the signs are changed to *r*_1_, *r*_2_ < 0, the empty strategy is promoted to SNE.

Supplementary Figure S2a explores the dynamics of an idealized Predator-Prey system with the interaction coefficients zeroed out, while keeping the intrinsic growth rates the same: i.e., *α* = 1.0, *β* = 0, *δ* = 0, *γ* = 1.5. The initial conditions were set at prey = 0.6, predator = 100. With no predatory interactions, the prey is free to grow exponentially while the predator’s population decays exponentially. Under this Nash equilibrium, frequency-dependent dynamics no longer drive the system dynamics. Nowak^3^ claims that replicator dynamics no longer adequately describe biological behaviors. To drive the point that only the imposed boundary condition drives the system dynamics, Supplementary Figure S2b zeros all model parameters: *α* = *β* = *δ* = *γ* = 0, and consequently *r*_1_ = *r*_2_ = 0. The deterministic dynamics no longer alter the population interaction terms. Under this condition, the system is governed by stochastic processes, such as the neutral drift described by Kimura^21^, until another deterministic process replaces frequency-dependent interactions, as we shall see in Supplementary Note S5.

Without a boundary condition, the idealized predator-prey model grows exponentially without bound. This is often accomplished by replacing the constant intrinsic growth coefficient with a function of the carrying capacity, *K*.

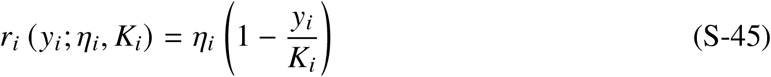

Mathematically, the notation *r*_*i*_ (*y*_*i*_; *η*_*i*_, *K*_*i*_) reads *r*_*i*_ is a function of *y*_*i*_ parameterized by *η*_*i*_ and *K*_*i*_. This function allows the terms in the parentheses to approach zero as the population density approaches *K*_*i*_. The parameterized carrying capacity function can now replace the constant growth coefficient.

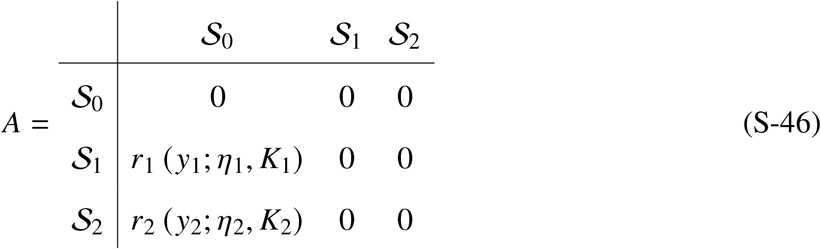

**Supplementary Figure S2:**
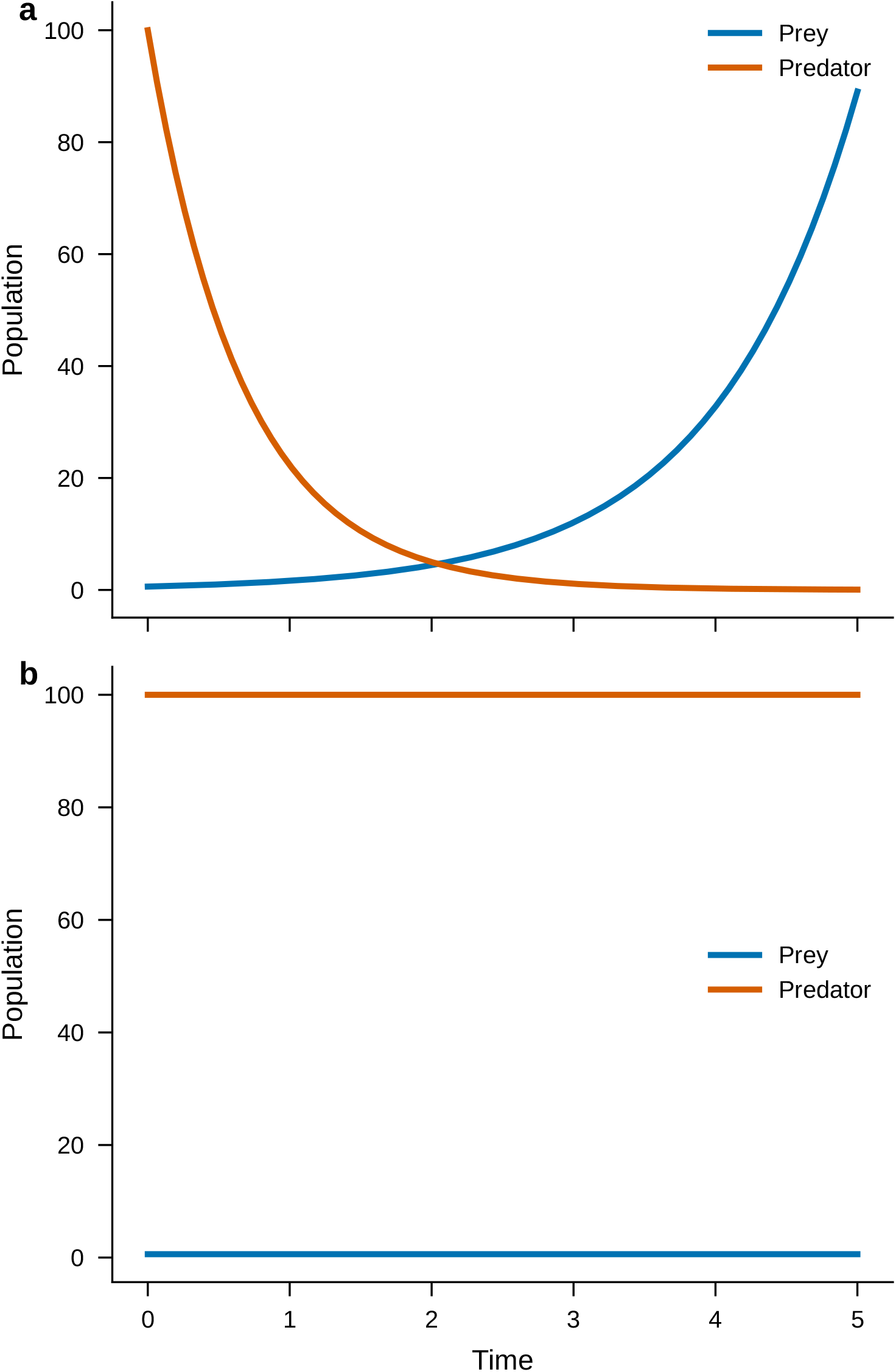
The ideal Predator-prey dynamics with initial conditions: prey = 0.6, predator = 100. a) Idealized parameters: *α* = 1.0, *β* = 0, *δ* = 0, *γ* = 1.5. b) No dynamics: *α* = *β* = *δ* = *γ* = 0.

Supplementary Note S5 uses this parametrized implementation of the boundary condition to formally derive the Pure Environmental Games.

## S5 The Pure Environmental Games

### The Pure Environmental Game

This work defines the Pure Environmental Games as any biological system in which the absolute population growth rate of the *i*-th member (*X*_*i*_) can be expressed as

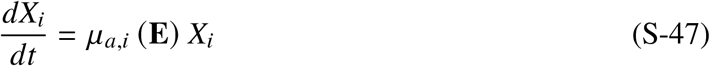

where

- **E** is the environmental vector specifying parameters such as pH, temperature, and concentration of any gaseous or aqueous chemicals. For engineered systems, **E** includes control parameters such as the hydraulic residence time, *θ* [*T* ].
- *μ*_*a,i*_ (**E**) the net specific accumulation rate of the *i*-th member [*T*^−1^]. Since **E** includes dynamic variables such as substrate concentration, *S*(*t*), the net specific accumulation rate *μ*_*a,i*_ (**E**) is implicitly time-dependent. Supplementary Note S6 will examine the behavior of the dynamic payoff matrix.

Following the derivation of the ideal LVM in Supplementary Note S4, the term *μ*_*a,i*_ (**E**) can be placed under the empty strategy column of the payoff matrix. The Pure Environmental Games redefine *S*_0_ and *x*_0_ as

- *S*_0_ - environmental strategy.
- *x*_0_ - environmental frequency.

When the frequency of the environmental strategy, *x*_0_, multiplies *μ*_*a,i*_ (**E**), it provides a mathematical framework to project the environmental dynamics onto the simplex, thereby exerting pressure on biological selection.

It should be noted that the term “pure” has two separate meanings. In replicator dynamics, each individual in a population comes with a hardwired phenotype representing a “pure” strategy. In the Pure Environmental Games, “pure” also refers to the payoff matrix structure: the entries *a*_*i j*_ = 0 for all *i, j* > 0, meaning biological strategies do not directly interact with each other. Instead, microorganisms receive payoffs from interacting with the environment, i.e., *a*_*i*0_. Later sections will show that *S*_*i*_ can represent microbial genera such as *Methanosarcina* and *Methanothrix*. The remainder of this section will derive a specific instance of the Pure Environmental Games, known as the idealized chemostat, from Environmental Biotechnology.

As we move from a mathematical model to a physical model, the use of internally consistent physical units becomes essential. This section describes physical dimensions using length (*L*), time (*T* ), mass (*M*), and energy (*E*) unless specified otherwise. The notation [·] should be read as “is in the units of.” The masses will be denoted by subscripts *M*_*X*_, *M*_*S*_, and *M*_*P*_ for the biomass, substrate, and product, respectively. For internal consistency, all masses will be expressed in terms of electron equivalents: 1 electron equivalent (or 1 eeq) is defined as 1 mole of electrons. The next section introduces all anabolic, catabolic, and decay reactions with the transfer of 1 eeq.

### Monod Kinetics

Microorganisms grow by taking up sources of energy and cellular building blocks from the surrounding environment. The Monod kinetics^40^ describes the growth rates of microorganisms limited by the substrate – a compound serving as the primary source of energy with a symbol *S* [*ML*^−3^]. The single-substrate Monod kinetics assumes that only one substrate (i.e., food or electron donor) is rate-limiting and has the form

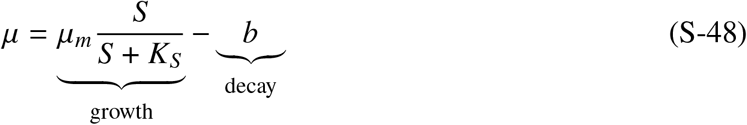

where

- *μ* - net specific growth rate [*T*^−1^]
- *μ*_*m*_ - maximum specific growth rate [*T*^−1^]
- *b* - endogenous decay rate [*T*^−1^]
- *S* - substrate concentration [*M*_*S*_ *L*^−3^]
- *K*_*S*_ - half-maximum rate substrate concentration [*M*_*S*_ *L*^−3^]

On the right-hand side, the first Monod term describes the generation rate of new biomass. The second term is the maintenance rate of microorganisms. For modeling environmental microorganisms, the Monod kinetics needs to be integrated with mass conservation relationships.

### The Idealized Chemostat

Environmental Biotechnology combines conservation laws, reaction kinetics, and reaction thermodynamics to model microbiological systems. The models can range from simple (e.g., reaction stoichiometry) to extraordinarily complex (e.g., 3-dimensional biofilm models). This work assumes the idealized chemostat model that microbial ecologists and engineers commonly use to develop microbial ecology principles and for designing wastewater treatment processes. The term “idealized” means that the system is homogeneous and the microorganisms are freely suspended without interactions (no flocs or biofilms). For a comprehensive treatment of the chemostats, the readers are referred to^13^.

The chemostat, also known as a Continuous Stirred-Tank Reactor (CSTR), is a microbiological reactor consisting of a continuous inflow, a continuous outflow, and a well-mixed growth chamber.

A combination of substrate, biomass, and product mass balance equations describes a chemostat:

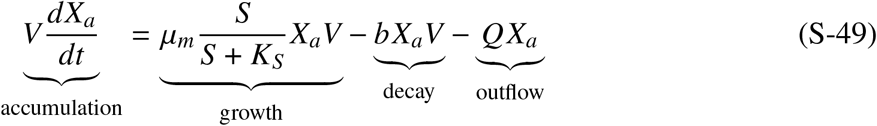

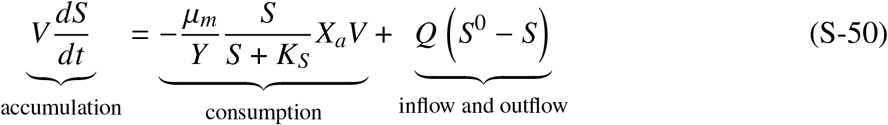

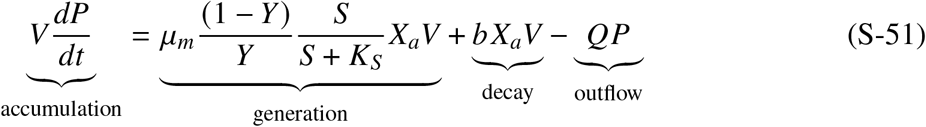

where

- *X*_*a*_ - active biomass concentration [*M*_*X*_ *L*^−3^]
- *Y* - true yield 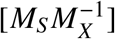
- *S*^0^ - influent substrate concentration [*M*_*S*_ *L*^−3^]
- *P* - effluent product concentration [*M*_*P*_ *L*^−3^]
- *V* - chemostat volume [*L*^3^]
- *Q* - hydraulic flow rate [*L*^3^*T*^−1^]

The subscripts *X* and *S* denote biomass and substrate, respectively.

To transform the mass-balance relationship, we define the hydraulic residence time (HRT) as *θ* [*T* ], which is also the inverse of dilution rate *D* [*T*^−1^]

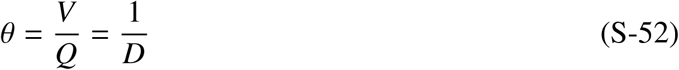

HRT is the average time a water parcel spends in a system. Another related fundamental parameter for Environmental Biotechnology is the solids residence time (SRT) or *θ*_*x*_, which Rittmann and McCarty ^13^ formally defines as

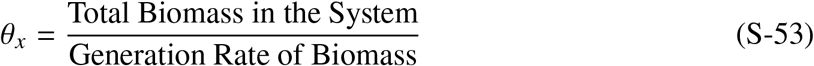

In wastewater engineering, SRT is also called the sludge age or the mean cell residence time. SRT is one of the most fundamental parameters in microbiological systems. SRT correlates with the structure and function of microbial communities in engineered and natural systems, including activated sludge^41^ and the human gastrointestinal tract^42^. When we apply this definition to the chemostat, we find that HRT=SRT:

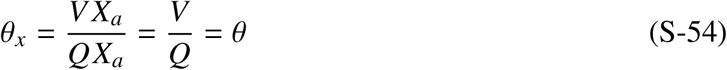

Using (S-52) and (S-54), the equations (S-49) and (S-50) can be simplified to:

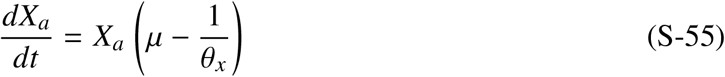

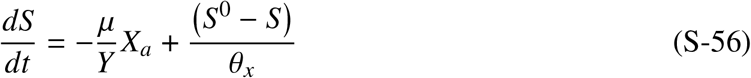

The modified biomass balance (S-55) shows structural similarity to the replicator equation (S-9). However, there are important distinctions. In the replicator equation, the strategy frequency increases when the individual strategy fitness exceeds that of the population average fitness. In microbiological systems, microbial biomass increases when the net specific rate of microorganisms exceeds the dilution rate. In environmental dynamics, microorganisms are racing against time (i.e., the term 1/*θ*_*x*_), and their growth rate is limited by the substrate (S-56). In the PEGs, the term in the parentheses (S-55) represents the net accumulation rate:

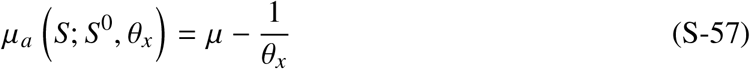

Following the projection (Supplementary Note S3), the biomass balance can be transformed into the replicator equation with the net accumulation appearing under the empty strategy.

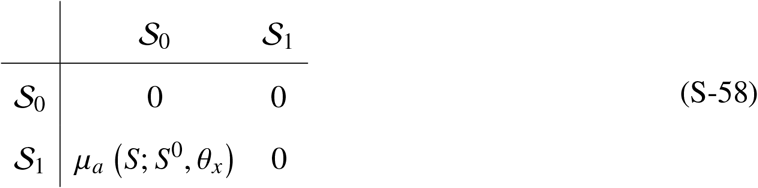

It should be noted that since the original chemostat system has one active biomass component, the corresponding replicator equation has two strategies: empty strategy and active-biomass strategy. *μ*_*a*_ is a function of the substrate concentration *S*, which is a dynamic variable. Therefore, to obtain the full dynamics of this environmental game, the biomass mass balance needs to be solved together with the substrate balance with *S*^0^ and *θ*_*x*_ as the boundary condition and control parameter, respectively.

### The Steady State

The steady-state occurs when the system dynamics stabilize, and system variables become time-invariant. For example, setting *dX*_*a*_/*dt* = 0 in (S-55) gives a fundamental relationship that holds at the steady-state

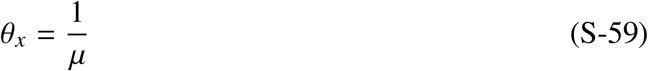

Rittmann and McCarty ^13^ call *θ*_*x*_ “a fundamental descriptor of the physiological status of the system” because it equals *μ*^−1^ at the steady state. Supplementary Note S6 explores how *θ*_*x*_ acts as a “lever” for environmental selection.

Remarkably, at steady state, *μ*_*a*_(*S*; *S*^0^, *θ*_*x*_) = 0, and the payoff matrix (S-58) simplifies to

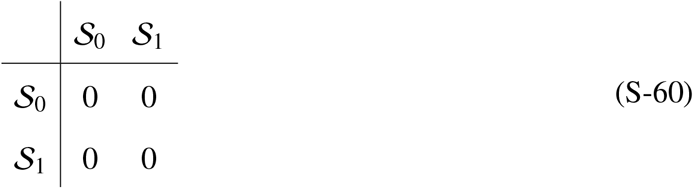

where all strategies, including the environmental strategy, become a Nash equilibrium. Does this mean that environmental conditions no longer exert any selective pressure across the simplex boundary?

To answer this question, it is helpful to examine the steady-state concentrations of substrate and biomass. Set the left-hand side (LHS) of (S-49) and (S-50) to zero, and divide the equations by *V*

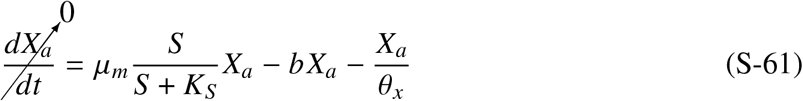

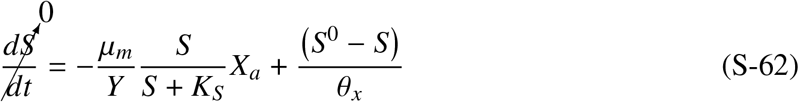

Then, rearranging the biomass balance gives the steady-state substrate concentration (*S*^∗^):

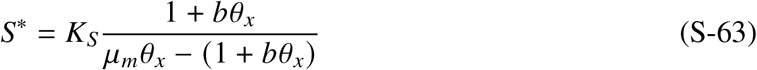

The steady-state active biomass concentration can be obtained by solving (S-61) for the first term on the right-hand side (RHS), substituting the results in (S-62), and solving for 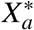:

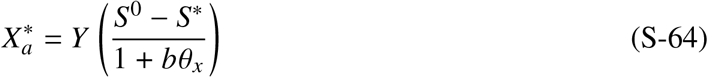

These algebraic solutions show two important features. First, the steady-state biomass concentration depends on the influent substrate concentration, *S*^0^, set at the boundary. When 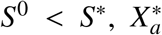 is negative, which is not physically possible and means that the chemostat is unable to support a steady state biomass (i.e., washout). Second, when *S*^0^ > *S*^∗^, *S*^∗^ and 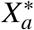 are stable dynamic equilibrium points – If *S* and *X*_*a*_ deviate from *S*^∗^ and 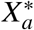, the payoff matrix becomes non-zero and drives the system back towards *S*^∗^ and 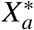.

This analysis differentiates a Nash equilibrium (a static equilibrium concept) from environmental pressures. At the steady state, it would be incorrect to interpret an absence of environmental pressure from *μ* (*S*; *S*^0^, *θ*_*x*_) = 0. The null payoff matrix indicates a stable dynamic equilibrium instead of environmental neutrality. When the system deviates from the steady state, *μ*_*a*_ (*S*; *S*^0^, *θ*_*x*_) ≠ 0 and the environmental pressures re-enter the payoff matrix, restoring the system towards the steady state. While this stability holds between the environmental strategy and biological strategies at steady state, Supplementary Note S6 explores neutrality among biological strategies under environmentally neutral selection.

### Limiting Steady State Values

There are two important limiting cases for subsequent sections. In the limit of a short *θ*_*x*_, minimal substrate is consumed (*S*^0^ ≈ *S*), and very little active biomass is generated. Then, (S-64) can be rearranged to define 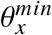:

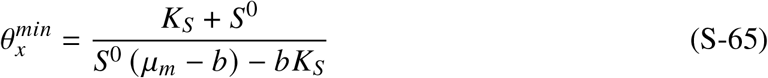

In the limit of a very large *S*^0^, (S-65) simplifies to define 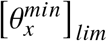,which is the absolute minimum time requirement for microbial growth^18^:

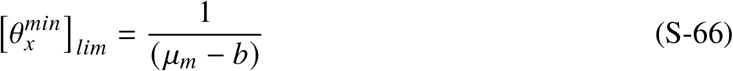

In the limit of a very large *θ*_*x*_, (S-63) simplifies to define 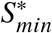, the minimum concentration capable of sustaining steady-state biomass^22^:

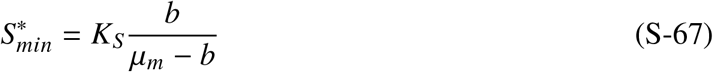

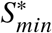 depends only on parameters related to microbial kinetics (i.e., *K*_*S*_, *μ*_*m*_, and *b*).

### The Generalized Pure Environmental Games

In the environment, microorganisms grow by taking up sources of energy and cellular components. In turn, they release waste products and modify the environment. A chain of microbial reactions forms a metabolic food web, in which the waste product of one microbe feeds another microbe. These metabolic interactions of microorganisms mediated through the environment can be described using the framework of multi-component reaction and transport modeling Steefel and MacQuarrie ^43^ . Here, we adopt the reaction framework to describe the generalized PEGs.

A multi-component model treats microorganisms and chemicals as components. When chemical and microbiological reactions consume components, the reaction stoichiometry can be written as

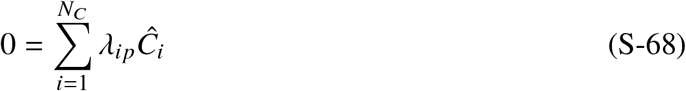

where *Ĉ*_*i*_ is the formula for component *i* [M] and *λ*_*i,p*_ is the stoichiometry coefficient for component *i* in a process *p* [*M*_*i*_ per reaction] when the process *p* is a reaction. *N*_*C*_ is the number of components. *λ*_*i,p*_ is negative for the reactant and positive for the product. For example, the chemostat example has consuming substrates and generating products. This growth reaction can be represented as

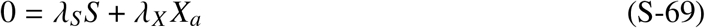

Then the true yield is

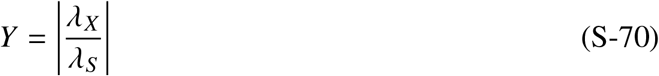

The conservation law for all components undergoing chemical and microbial reactions can be written as

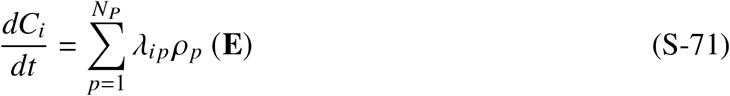

where *ρ*_*p*_ is the rate of *p*-th process and *N*_*P*_ is the number of processes.

The conservation law is consistent with the generalized PEGs if it can be rewritten in the form

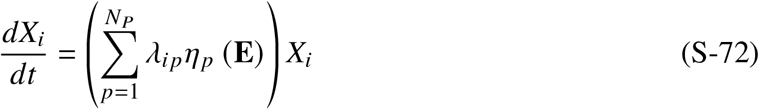

where *η*_*p*_ is the specific rate of *p*-th process and *N*_*P*_ is the number of processes. The process rates, *η*_*p*_, are positive by convention. Comparing this equation to the definition of the pure environmental game (S-47) gives

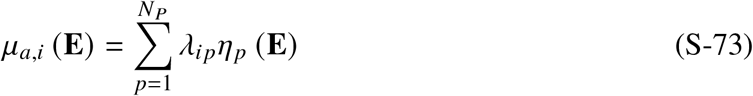

The chemostat example (S-49) illustrates this where

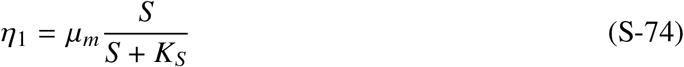

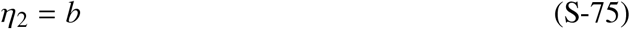

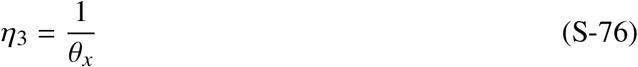

with the stoichiometry coefficients *λ*_*a*1_ = 1, *λ*_*a*2_ = −1, and *λ*_*a*3_ = −1 Note that the first two are microbiological processes, while the third is a physical transport process.

Table S2 summarizes the one-species chemostat model in a format commonly known as the Gujer matrix. In the Gujer matrix, each row represents a process. When the process is a reaction, it is described by (S-68). For transport processes, influent is positive and effluent is negative. The chemostat example in this section had six processes: growth, decay, biomass outflow, substrate inflow, substrate outflow, and product outflow. The middle section holds the stoichiometry coefficients, *λ*_*p,i*_, noting that the indices i and p are transposed relative to (S-71). The middle section is also known as the stoichiometry matrix, **Λ**. The final column shows the process rates, *ρ* (**E**). The Gujer matrix serves as the recipe for the conservation law, which can be restored by matrix multiplication.

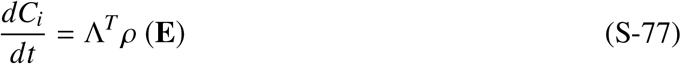

**Supplementary Table S2:**
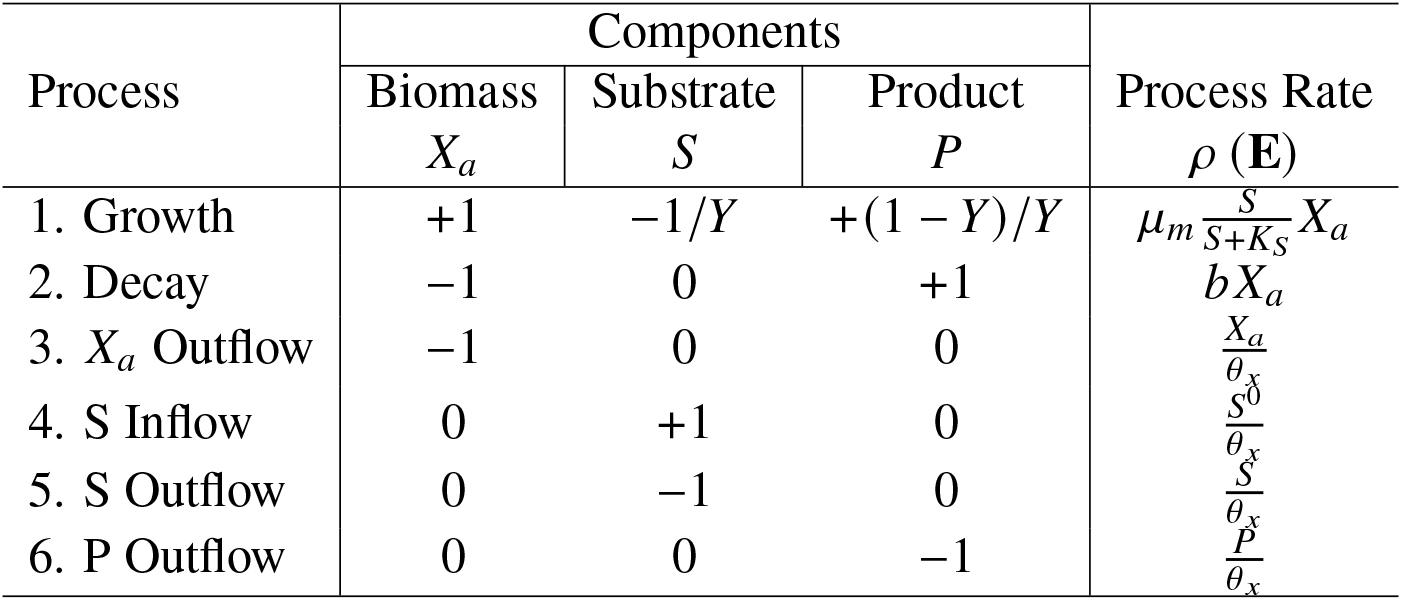
Gujer matrix for the one-species chemostat model.

### Energy Dissipation Rate

The second law of thermodynamics states that the total entropy of an isolated system must increase, Δ*S* > 0. A dilute aqueous system approximates the isothermal condition due to water’s high heat capacity, and the Gibbs free energy provides an equivalent statement of the second law. For a biological reaction index by *p*, the Gibbs free energy of reaction, Δ*G*_*p*_ [*E M*^−1^], can be written as

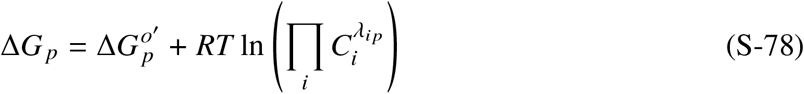

where 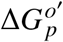 is the standard Gibbs free energy of the reaction [*E M*^−1^], *R* is the ideal gas constant (8.314 J/K-mol), and *T* is the absolute temperature in Kelvin. Here, the superscript “o” denotes standard condition and the prime denotes pH=7. Δ*G*_*p*_ must be less than zero for the reaction to occur spontaneously. 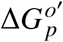 is determined from the standard Gibbs free energy of formation [*E M*^−1^] for each component in the reaction multiplied by the stoichiometry

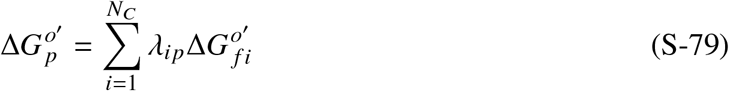

For simplicity, assume that the energy dissipated by microorganisms is transported out of the system by the bulk fluid. The energy dissipation rate of the system, 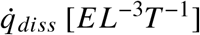, can be written as the product of the process rate and the Gibbs free energy of the overall reaction.

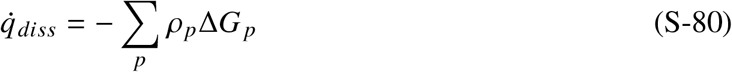

The system will perform a set of reactions that are spontaneous overall (exergonic), and the dissipation rate will be positive. During the synthesis of new biomass, microorganisms partition electrons into cell synthesis and energy generation. When Y has the units of 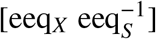, it is numerically equivalent to the symbol 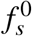 used in Rittmann and McCarty ^13^ . Since the microorganisms perform anabolic and catabolic reactions in parallel, it is convenient to write the overall process as the sum of anabolic and catabolic reactions (*R*_*p,ana*_ and *R*_*p,cat*_ respectively):

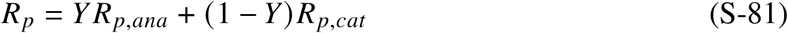

Here, *R*_*p*_, *R*_*p,ana*_, and *R*_*p,cat*_ all assume 1 mole of electrons (or 1 electron equivalent = 1 eeq) transferred. However, *ρ*_*p*_ are written in terms of the rate of biomass generation. Therefore, we define *σ*_*p,ana*_ = 1 and *σ*_*p,cat*_ = (1 − *Y* )/*Y* . Supplementary Table S3 summarizes the energy dissipation matrix for the one-species chemostat model. Here, both **Σ** and Δ**G** are *N*_*P*_ × 2 matrices. For the endogenous decay processes, there is no cell synthesis and *σ*_*p,ana*_ = 0. Both *σ*_*p,ana*_ = *σ*_*p,cat*_ = 0 for transport, as they are not reactions. The overall reaction can be divided into anabolic and catabolic reactions

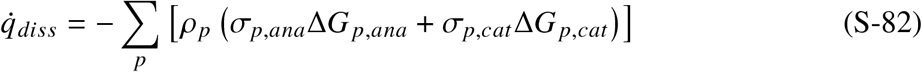

Or more generally, it can be calculated via matrix operations

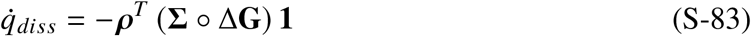

where ° denotes element-wise (Hadamard) product. For the chemostat system, the energy dissipation rate is written as

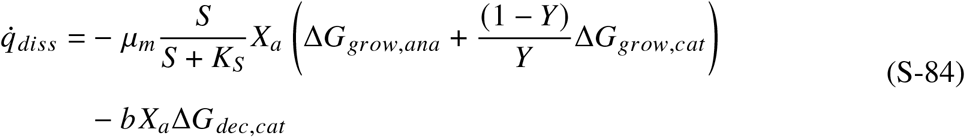

Finally, the specific energy dissipation rate of microorganisms, 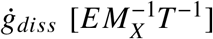, is defined by dividing this equation by *X*_*a*_,

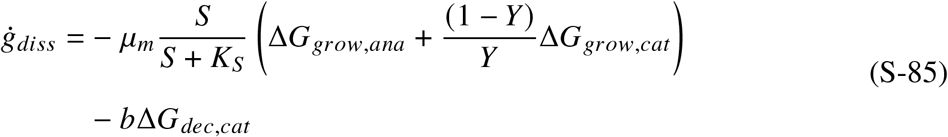

**Supplementary Table S3:** Energy dissipation matrix for the one-species chemostat model.

| Process | $\Sigma$ | | $\Delta G$ | | $\rho(\mathbf{E})$ |
| --- | --- | --- | --- | --- | --- |
| | $\sigma_{p,ana}$ | $\sigma_{p,cat}$ | $\Delta G_{p,ana}$ | $\Delta G_{p,cat}$ | |
| 1. Growth | 1 | $(1 - Y)/Y$ | $\Delta G_{grow,ana}$ | $\Delta G_{grow,cat}$ | $\mu_m \frac{S}{S+K_S} X_a$ |
| 2. Decay | 0 | 1 | 0 | $\Delta G_{dec,cat}$ | $b X_a$ |
| 3. $X_a$ Outflow | 0 | 0 | 0 | 0 | $\frac{X_a}{\theta_x}$ |
| 4. S Inflow | 0 | 0 | 0 | 0 | $\frac{S^0}{\theta_x}$ |
| 5. S Outflow | 0 | 0 | 0 | 0 | $\frac{S}{\theta_x}$ |
| 6. P Outflow | 0 | 0 | 0 | 0 | $\frac{P}{\theta_x}$ |

## S6 Environmental Selection Dynamics

The PEGs provide a framework to describe environmental games operating in parallel to the neutral selection game. This section explores the r/K selection game as an instance of PEGs with *Methanosarcina* and *Methanothrix* (formerly *Methanosaeta*) as two classic microorganisms competing for a shared resource – acetate in this case^15,44^. Six dynamic regimes are identified by exploring the parameter space of the r/K selection game.

Extended Data Table 2 summarizes the microbial kinetic parameters for *Methanothrix* and *Methanosarcina. Methanosarcina*’s competitive advantage to grow rapidly is characterized by the higher *μ*_*m*_ (r-strategy). *Methanothrix*’s competitive advantage to scavenge the scarce acetate is characterized by the lower *K*_*S*_ (K-strategy).

### Anticipated Steady State

The steady-state substrate concentration, *S*^∗^, (S-63) was developed independently by two separate traditions. For engineers, *S*^∗^ represents the effluent quality and water quality standard they must meet to avoid compliance violations and expensive penalties^18^. For microbial ecologists, *S*^∗^ (or more commonly *R*^∗^) represents the anticipated substrate concentration microorganisms will “experience” when the system reaches the steady-state and provides a basis for competitive exclusion^19,20^. *θ*_*x*_ is the “lever” that engineers have to control water quality, and microbial ecologists have to control microbial community composition.

Supplementary Figure S3 summarizes the relationship between *S*^∗^ and *θ*_*x*_ using (S-63), which represents outcomes if *Methanosarcina* and *Methanothrix* were grown in separate chemostats. As shown in (S-66), 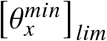 is the minimum time requirement for growth. When 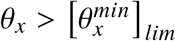, the methanogens are able to grow and *S*^∗^ increases with 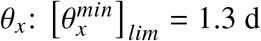 for *Methanosarcina* and 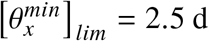 for *Methanothrix*, respectively.

**Supplementary Figure S3:**
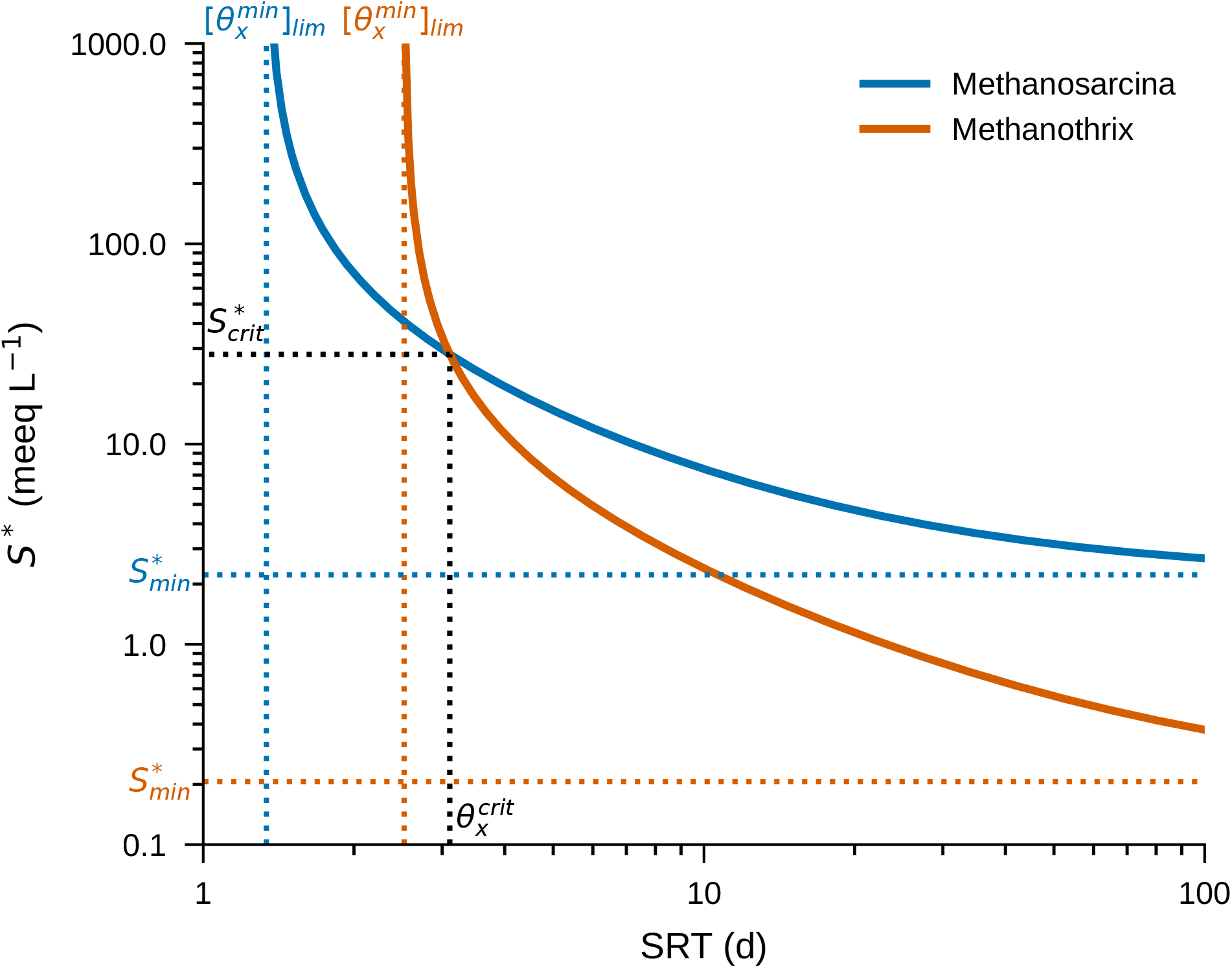
The relationship between the anticipated steady-state substrate concentration, *S*^∗^, and the solids residence time, *θ*_*x*_, if *Methanosarcina* and *Methanothrix* were grown in separate chemostats. Parameters are summarized in Table 2.

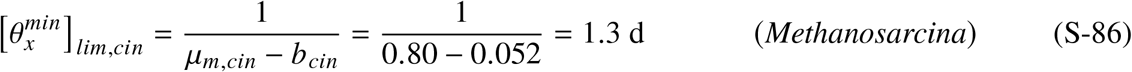

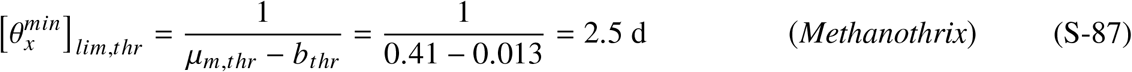

Furthermore, *S*^∗^ is a monotonically decreasing function of *θ*_*x*_ and approaches the value of 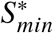 as defined in (S-67).

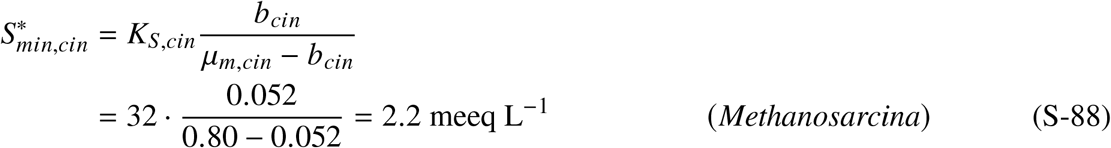

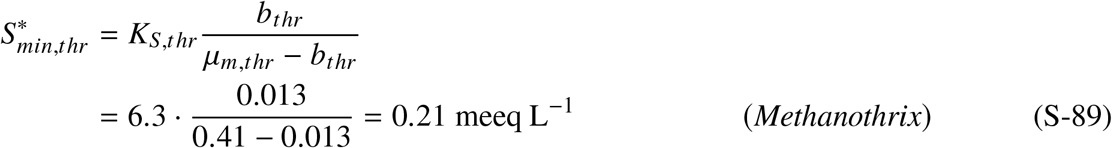

The 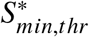 of 0.2 meeq L^−1^ or (1.7 mgCOD L^−1^) is several factors lower than a range of 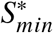 estimates 5.8 to 31.3 mgCOD L^−1^ obtained by McCarty and Bae ^45^ likely because of thermodynamic inhibition. Since McCarty and Bae ^45^ estimates *K*_*Ac*_ between 106 to 577 mgCOD L^−1^ (or 13 to 72 meeq L^−1^) for the same digesters, this work considers thermodynamic inhibition by an upward correction of *K*_*S,thr*_ from 6.3 to 22 meeq L^−1^ only for Regime V and VI (Extended Data Table 2). Therefore, the updated 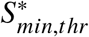 value for *Methanothrix* is

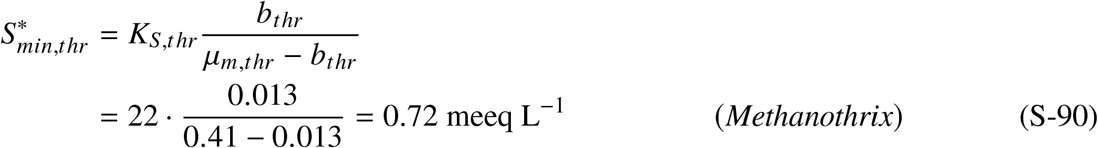

For calculating the specific energy dissipation by *Methanosarcina* and *Methanothrix*, Δ*G*_*ana*_ needs to be calculated at the steady state. Extended Data Table 3 summarizes the reaction stoichiometry for the anabolic reaction (cell synthesis), catabolic reaction (energy generation), and decay reactions for acetoclastic methanogens. Assuming pH=7, pCO2 = 0.5 atm, [NH_4_]=0.01 mmol/L, and unit activity for biomass and water, Δ*G* value can be expressed as a function of *S*_*Ac*_

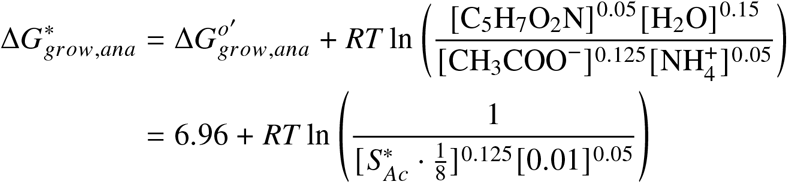

Then, at 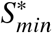 the energy values for Δ*G*_*grow,ana*_ are

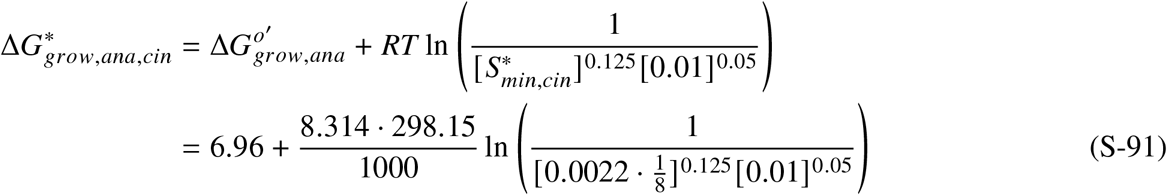

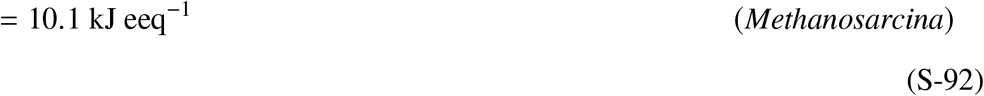

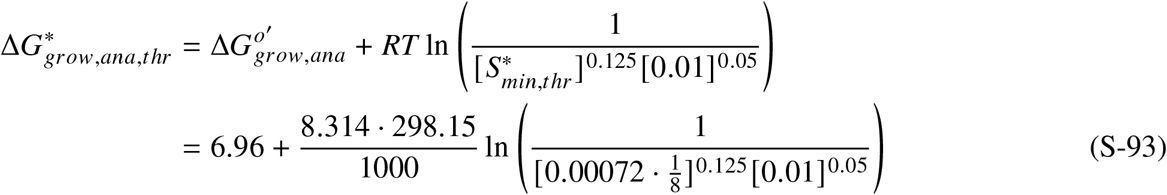

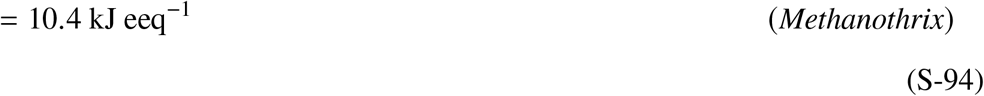

Similarly, Δ*G*_*grow,cat*_ at 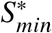 are

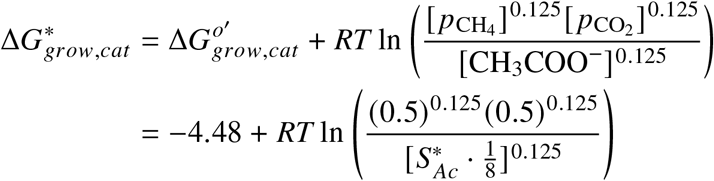

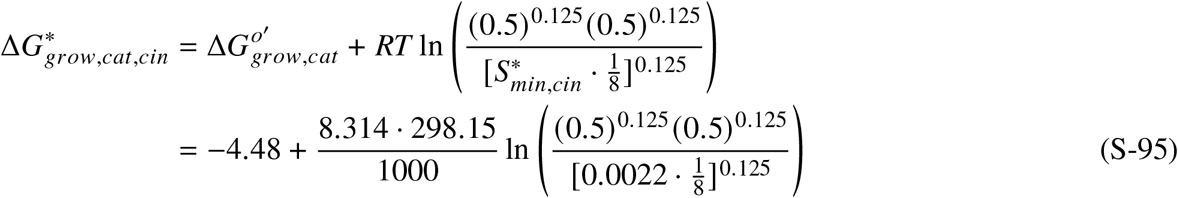

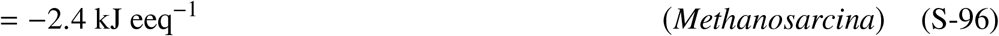

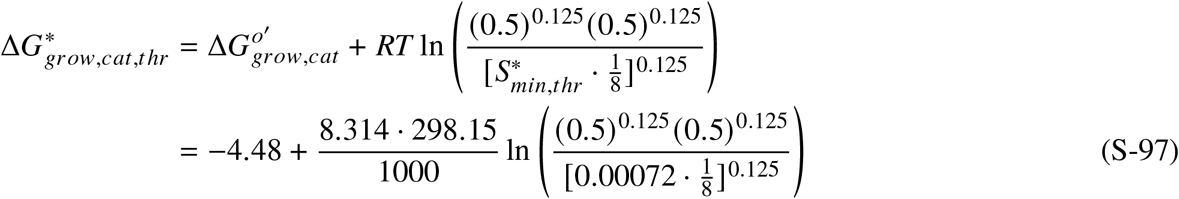

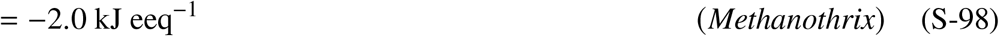

Finally, Δ*G*_*dec,cat*_ at 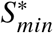 for both *Methanosarcina* and *Methanothrix* are

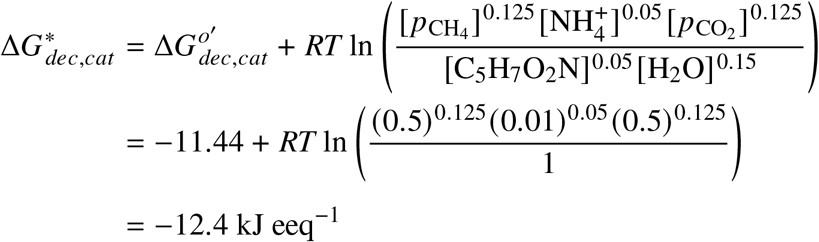

Then, the anticipated specific energy dissipation rate can be calculated at 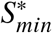 using (S-85).

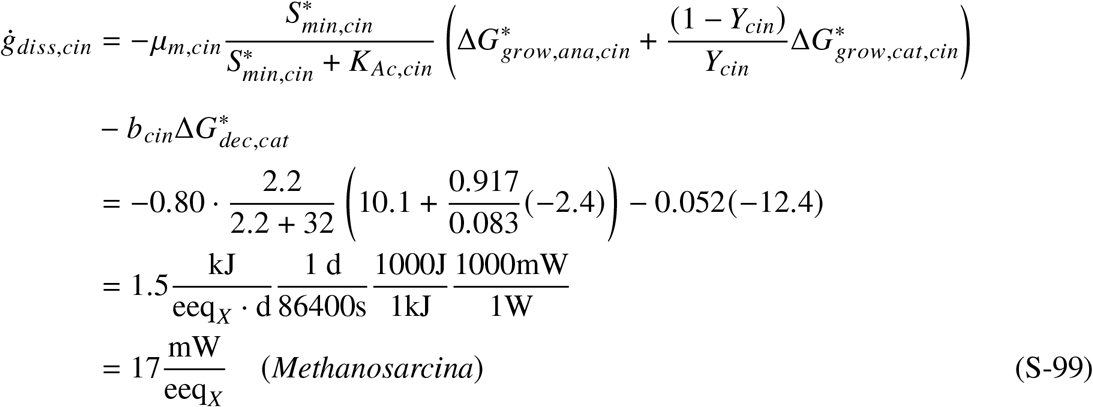

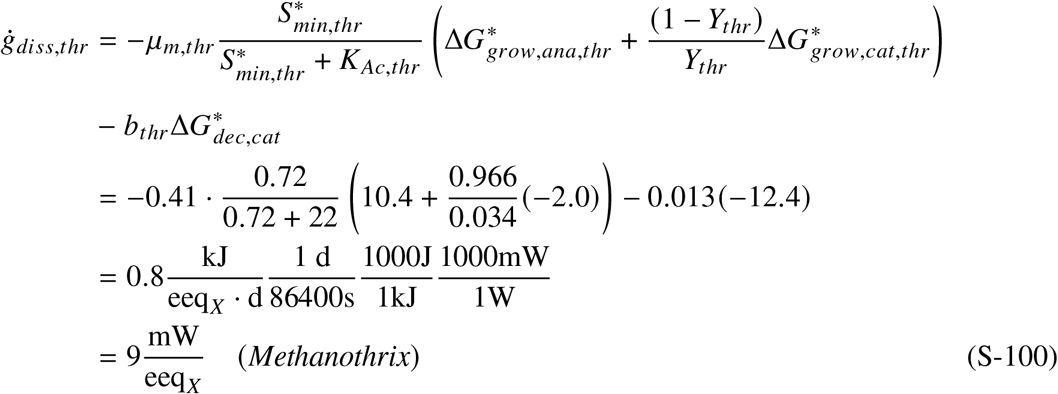

Therefore, the specific energy dissipation rates of maintaining one electron equivalent of *Methanosarcina* and *Methanothrix* are 17 and 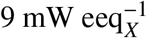 at 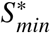, respectively.

In Supplementary Figure S3, there is a point where the lines for *Methanothrix* and *Methanosarcina* cross. This can be called the critical point where the dominant community member shifts, as the next subsection shows. At the critical point, *Methanosarcina*’s ability to grow rapidly is exactly neutralized by *Methanothrix*’s ability to scavenge acetate. For this *Methanothrix*-*Methanosarcina* system, the critical point occurs at 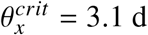 with 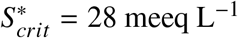.

### Dynamics of *Methanosarcina-Methanothrix* System

The anticipated steady states and critical points established in the previous subsection provide anchors for interpreting system dynamics. A system of equations – (S-101), (S-102), and (S-103) – describes the *Methanosarcina*-*Methanothrix* system

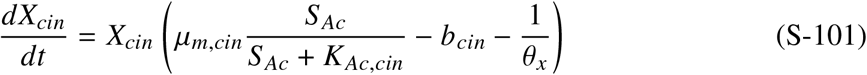

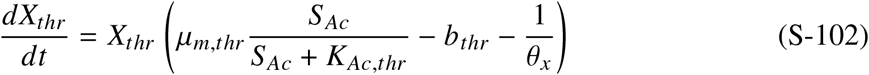

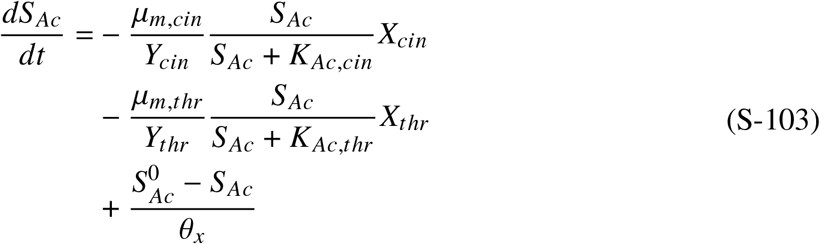

Describing the energy dissipation rate requires integration of reaction stoichiometry with reaction kinetics. Table 3 summarizes the anabolic, catabolic, and decay reactions for acetoclastic methanogens. All acetoclastic methanogens share the same Δ*G* values for the anabolic, catabolic, and endogenous decay reactions. However, the coefficient *Y*_*i*_ combines Δ*G* values in different proportions. Therefore, the energy dissipation rate for the chemostat following (S-83) is

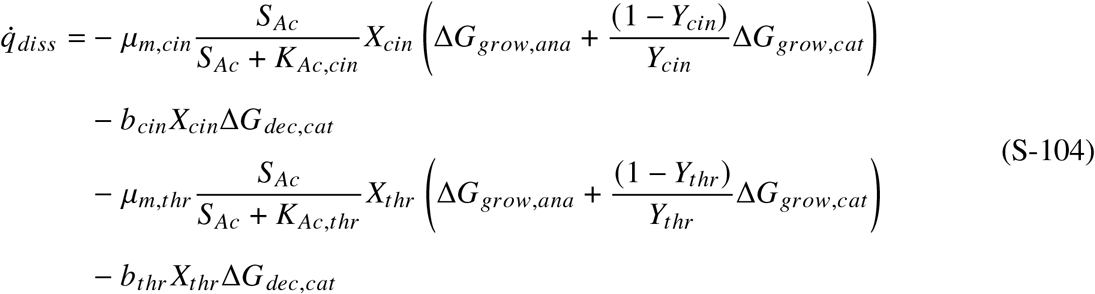

Extended Data Table 1 summarizes the Gujer matrix for the *Methanosarcina*-*Methanothrix* chemostat model.

### Washout 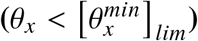 (Regime I)

Consider an idealized chemostat with a highly favorable 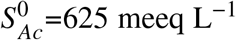 and initial *X*_*cin*_ = 500 meeq L^−1^ and *X*_*thr*_ = 500 meeq L^−1^. Extended Data Fig. 1 shows the *Methanosarcina*-*Methanothrix* dynamics in the absolute coordinate with 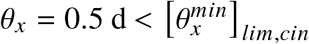 and 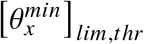. Initially, the acetate concentration sharply decreases from *S*_*Ac*_ = 625 to 1.6 meeq L^−1^ within 2.5 h due to the high concentration of *Methanosarcina* and *Methanothrix* (Extended Data Fig. 1a). Correspondingly, 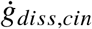 drops from 340 to 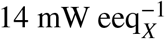 and 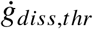 from 510 to 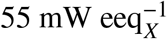. The energy dissipation rate for *Methanothrix* is higher than *Methanosarcina* because it requires a higher proportion of catabolic reaction per unit of biomass (i.e., 1 − *Y*_*thr*_ > 1 − *Y*_*cin*_). The decrease in biomass concentration is more gradual as the physical biomass removal lags behind microbiological acetate consumption. During the first day (*t* < 1 d), the biomass concentration for *Methanosarcina* decreases from 500 to 76 meeq L^−1^ and *Methanothrix* decreases from 500 to 82meeq L^−1^. *Methanothrix* fares marginally better as a K-strategist by scavenging acetate at a low concentration. After *t* = 1 d, both populations are washed out, and *S*_*Ac*_ returns to the influent level by Day 8. In tandem, the energy dissipation rates return to the highest levels (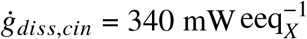 and 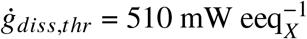), but are not sufficient to let microorganisms grow fast enough (Extended Data Fig. 1c).

Extended Data Fig. 1b shows the same dynamics projected onto the simplex. Initially, the simplex is fully occupied by an equal amount of *Methanosarcina* and *Methanothrix* strategies. During the first day (*t* < 1 d), *Methanothrix* gains in simplex frequency from 0.5 to 0.52, while *Methanosarcina* loses in simplex frequency from 0.5 to 0.48. Thereafter, *Methanothrix* is washed out, but between Days 1 and 3 *Methanosarcina* momentarily gains in simplex frequency from 0.48 to 0.53 by taking advantage of the elevated *S*_*Ac*_ as an r-strategist. However, this resistance proves futile. Both methanogen strategies are washed out, yielding the simplex to the environmental strategy.

Why does the environmental strategy resist invasion? At time zero, the net-specific accumulation rates for *Methanosarcina* and *Methanothrix* can be calculated by examining the terms in the parentheses for (S-101) and (S-102) at *t* = 0 d:

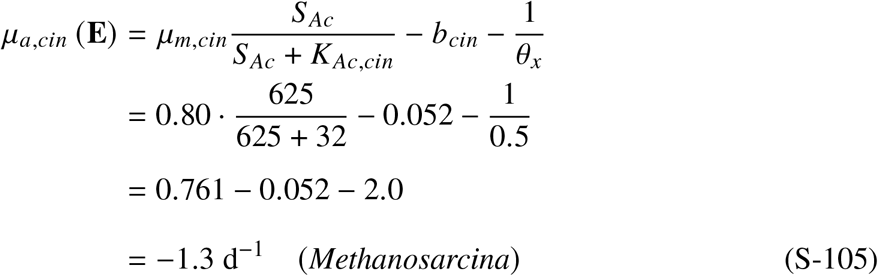

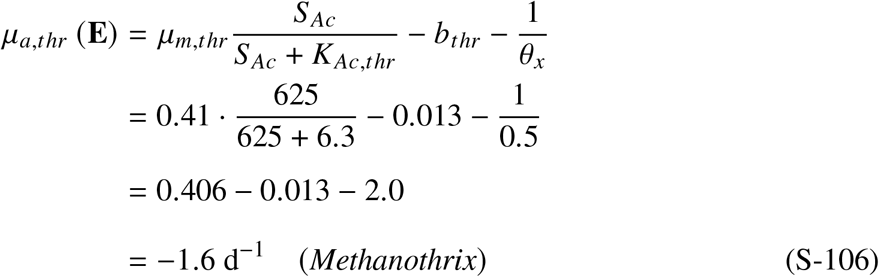

Using these values, the dynamic payoff matrix at t=0 (S-107) reveals that the environmental strategy is SNE (Supplementary Notes S2 and S4). This washout case 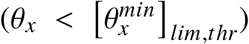, formally described by Lawrence and McCarty ^18^, prohibits microorganisms from growing in a chemostat. This work shows that this condition is equivalent to the scenario where the environmental strategy is an SNE. If environmental conditions do not provide enough time for microbes to grow, life cannot emerge.

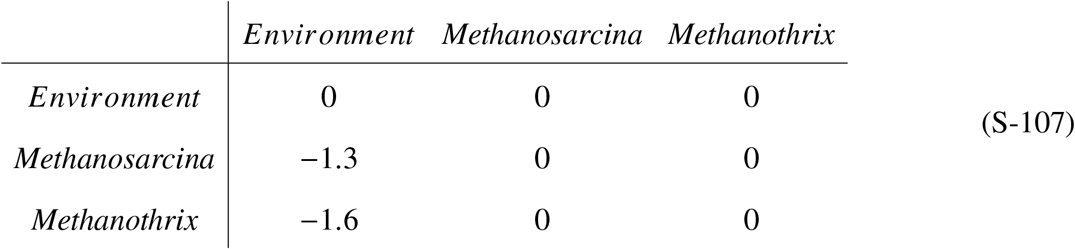

### r-dominates 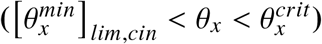 (Regime II)

The washout (Regime I) example illustrates that time can prohibit microorganisms from invading the environment. This example increases *θ*_*x*_ = 2 d, such that 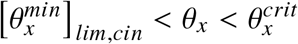 to illustrate Regime II: r-dominates. The final state of the Regime I simulation was taken to be the initial condition: *X*_*cin*_ = 5.7 × 10^−4^ meeq L^−1^ and *X*_*thr*_ = 4.2 × 10^−5^ meeq L^−1^. The influent acetate is kept at 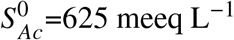.

Extended Data Fig. 2a shows the *Methanosarcina*-*Methanothrix* dynamics in the absolute coordinate. At *t* = 0 d, the net specific accumulation rates for *Methanosarcina* and *Methanothrix* according to (S-101) and (S-102) are

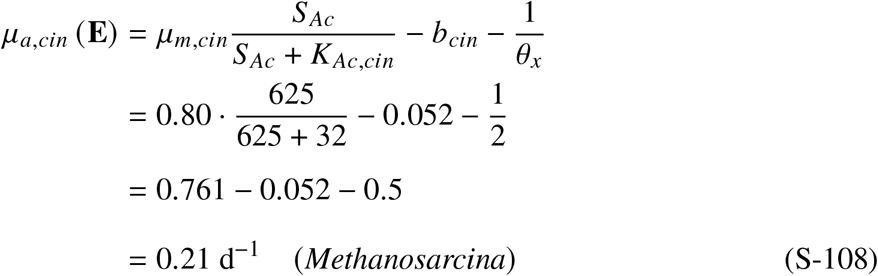

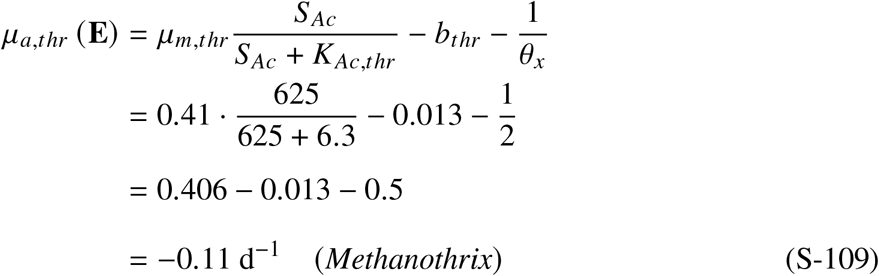

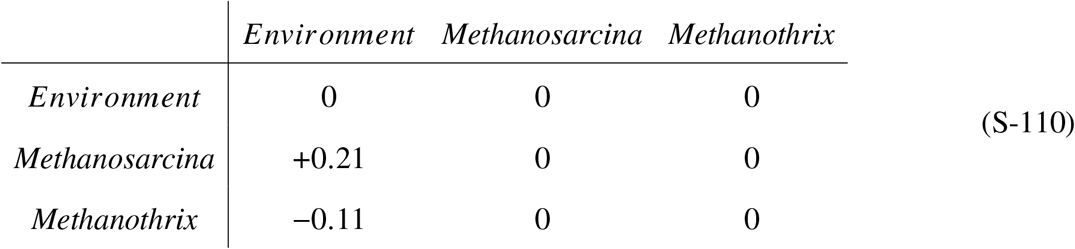

Using these values, the dynamic payoff matrix at t=0 (S-110) reveals that the positive net accumulation rate of *Methanosarcina, μ*_*a,cin*_ (**E**) = +0.21, disqualifies the environmental strategy from being SNE and ESS. On the other hand, *Methanosarcina* receives positive payoff from the environment to gain abundance from *X*_*cin*_ = 5.7 × 10^−4^ meeq L^−1^ at *t* = 0 d to the steady-state level of *X*_*cin*_ = 42 meeq L^−1^ by *t* = 40 d. In tandem, the steady-state acetate concentration drops to *S*_*Ac*_ = 71 meeq L^−1^ and *Methanothrix* is being washed out (*X*_*thr*_ = 1.4 × 10^−8^ meeq L^−1^ at *t* = 40), unable to receive positive payoff from the environment. Correspondingly, 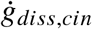 drops from 340 to 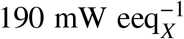 and 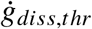 from 510 to 390 mW 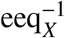 (Extended Data Fig. 2c). Therefore, *Methanosarcina* can accumulate and sustain at the same or lower energy dissipation rates given sufficient time at *θ*_*x*_= 2 d. These observations are corroborated by displacement of *Methanothrix* by *Methanosarcina* in the simplex (Extended Data Fig. 2b). Using these values, the net specific accumulation rates for the dynamic payoff matrix at *t* = 40 d are

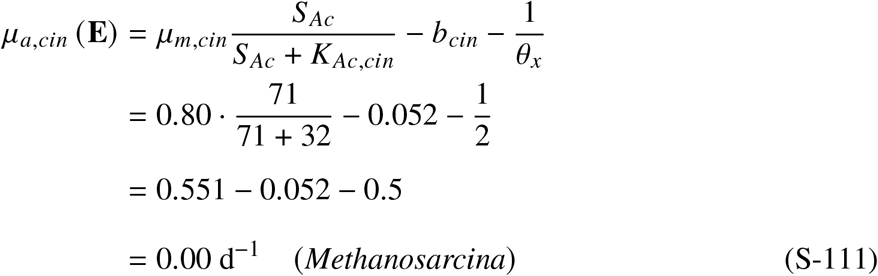

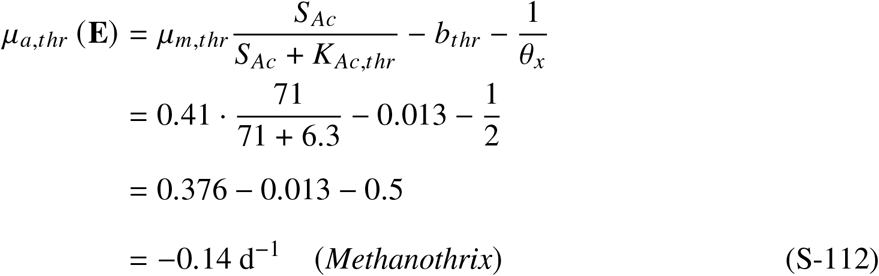

At the steady-state, *μ*_*a,cin*_ (**E**) = 0.00 (S-113). Therefore, using these values, the dynamic payoff matrix reveals that the environmental strategy is promoted to a Nash equilibrium at the steady state. While both *Methanosarcina* and environmental strategies are Nash equilibrium strategies, they have deterministic outcomes because environmental dynamics drive these strategies towards a steady state. Therefore, the steady-state is not environmentally neutral.

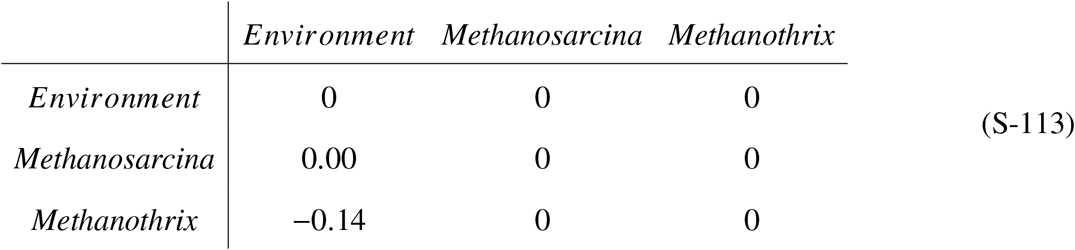

### K-dominates 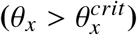 (Regime IV)

This example uses the final state of Regime II as the initial condition: *X*_*cin*_ = 42 meeq L^−1^, *X*_*thr*_ = 1.4 × 10^−8^ meeq L^−1^, *S*_*Ac*_ = 71 meeq L^−1^. As stated earlier, this *Methanosarcina*-*Methanothrix* system has a 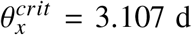 (four significant digits are shown here for illustrative purposes). This example adds a tiny *δ* = 0.01 d to this critical value and sets *θ*_*x*_ = 3.107 + 0.01 = 3.117 d to illustrate the profound effects of a minuscule deviation.

Extended Data Fig. 3a shows the *Methanosarcina*-*Methanothrix* dynamics in the absolute coordinates. Upon changing the *θ*_*x*_, the system takes about 100 days to reach a new stable state of *S*_*Ac*_ = 27.9 meeq L^−1^. In tandem, 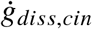 drops from 190 to 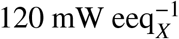 and 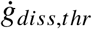 from 390 to 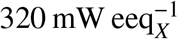 (Extended Data Fig. 3c). The net specific accumulation rates for *Methanosarcina* and *Methanothrix* are negligibly small:

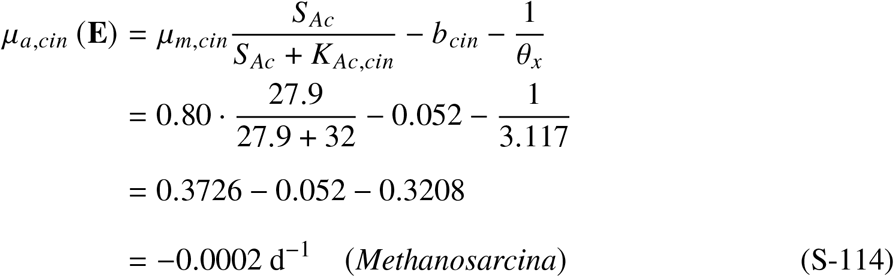

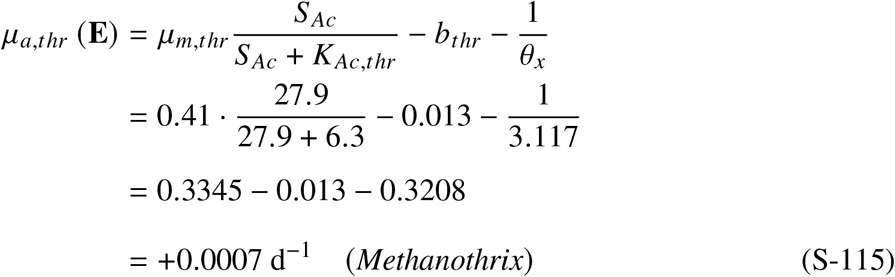

Placing these *μ*_*a*_ values at t = 100 d in the dynamic payoff gives a false appearance that the system is at an environmentally neutral selection (see Regime III. Environmental Neutrality).

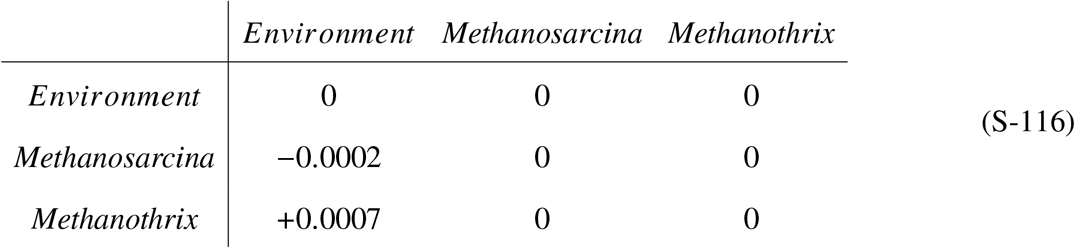

For the first 20,000 days, or 55 years, there is no apparent change in *X*_*cin*_ and *S*_*Ac*_. However, quietly *Methanothrix* grows exponentially from 1.4×10^−8^ meeq L^−1^ to *X*_*thr*_ = 0.078 meeq L^−1^. Exponential growth continues beyond this point, as *Methanothrix* becomes the dominant methanogen, displacing *Methanosarcina* by 30,000 days. Therefore, despite having a higher specific energy dissipation rate, *Methanothrix* can come to dominance by having a lower *K*_*S*_. It takes a total of 82 years for *Methanothrix* to reach steady state. Extended Data Fig. 3b shows that this exponential growth of *Methanothrix* has continued for the entire 82 years!

This example illustrates that in an idealized chemostat system, a minuscule competitive advantage derived from having 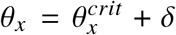 can compound over the years and allow rare species to become dominant. While maintaining an idealized chemostat with a tiny delta is not practical for engineered systems, 82 years is only a fraction of a second on an evolutionary scale.

### Environmentally Neutral Selection 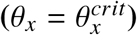 (Regime III)

Even a tiny deviation from 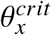 allows the r/K strategies to govern environmental dynamics in an idealized chemostat. However, when 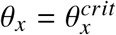 (which is not practical to simulate numerically), environmentally neutral selection challenges this idea. At 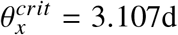 with 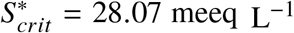 (four significant digits are shown here for illustrative purposes), the net specific accumulation rates of *Methanosarcina* and *Methanothrix* both equal zero:

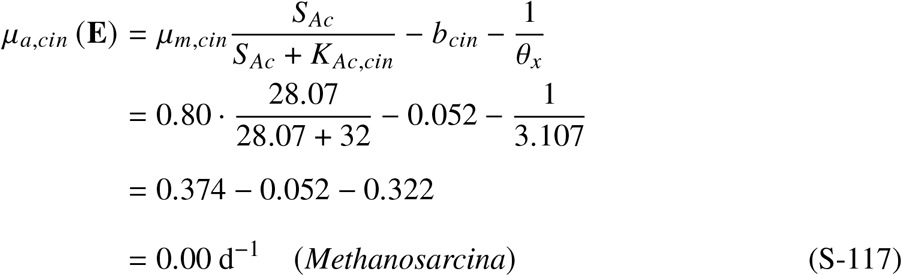

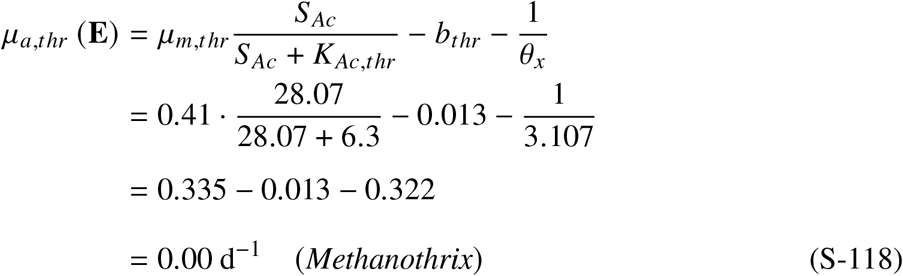

When these values are placed in the dynamic payoff matrix, all entries are zero (S-119).

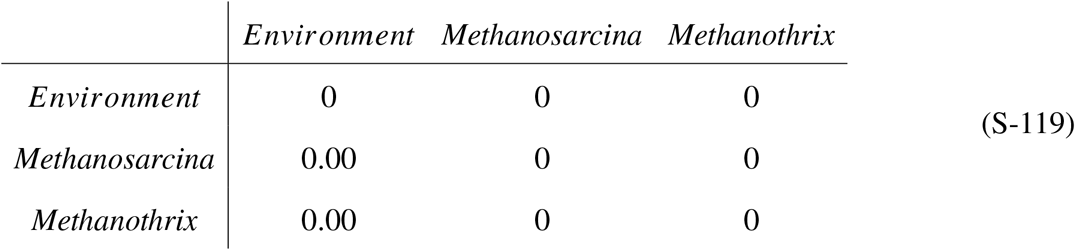

Therefore, *Methanosarcina* and *Methanothrix* have equal fitness. Following Nowak ^3^, who calls frequency-neutral interactions between Nash equilibrium strategies neutral selection, we define the analogous condition in the environmental game as environmentally neutral selection with caveats: In the traditional game, the condition *a* = *c* and *b* = *d* defines neutral selection. In the r/K selection game, 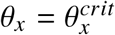 and *S*^0^ > *S*^∗^ ensure an environmentally neutral selection, while equal fitness is an outcome of the steady state.

While 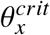 is a useful theoretical construct, the probability of a system operating at exactly this point in an idealized chemostat system is vanishingly small. Near 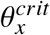, deterministic environmental selection cannot predict the outcome in an idealized system such as the chemostat. Mechanistic coexistence mechanisms – such as spatial heterogeneity in flocs^46^ and time-varying environments in sequencing batch reactors^15^ – may obscure this boundary in non-idealized systems. In the absence of such mechanisms, stochastic drift in the spirit of Kimura ^21^,^47^ determines which species ultimately prevails.

### 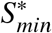 **(Regime V)**

This example uses the final state of Regime IV as the initial condition: *X*_*cin*_ = 0.034 meeq L^−1^, *X*_*thr*_ = 20 meeq L^−1^, and 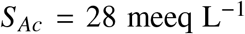. The influent is kept at 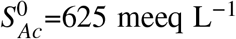. The solids residence time is adjusted to *θ*_*x*_ = 100, 000 d or 274 years to simulate severe mass-transfer resistance for the chemostat.

Extended Data Fig. 4a shows the *Methanosarcina*-*Methanothrix* dynamics in the absolute coordinates. Upon startup, *S*_*Ac*_ immediately drops from 28 to 0.0021 meeq L^−1^ due to severe mass transfer resistance. During this 1000 d period, Extended Data Fig. 4b shows that *Methanothrix* had been decaying. Thereafter, the system slowly relaxes to the final state of *X*_*thr*_ = 0.0163 meeq L^−1^ and *S*_*Ac*_ = 0.72 meeq L^−1^, while *Methanosarcina* washes out.

The final *X*_*thr*_ = 0.0163 meeq L^−1^ is consistent with the *S*_*min*_ = 0.72 meeq L^−1^ (S-89) which is realized at a large extreme of *θ*_*x*_. At this limit, the final biomass concentration can be calculated by replacing *S* with 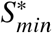, the steady-state biomass equation (S-64)

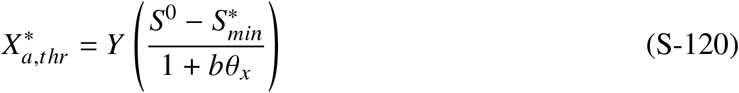

For example, the predicted steady-state biomass concentration for *Methanothrix* with *θ*_*x*_ = 100, 000 d and 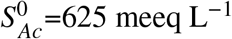.

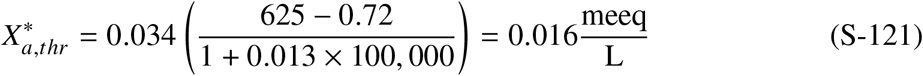

which is equal to the dynamic simulation final state. In fact, the steady-state biomass concentration will approach zero as *θ*_*x*_ approaches infinity, meaning the steady-state biomass approaches a vanishingly small but nonzero concentration.

### *Persistence*. (Regime VI)

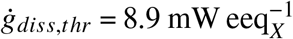 represents the true minimum amount of energy dissipation required for *Methanothrix* to invade the environment (Extended Data Fig. 4c). Using 2.8 ×10^−13^ g per cell^13^ and 113 g of cell per 20 eeq_*X*_ according to the cellular formula C_5_H_7_O_2_N in Extended Data Table 3, this equates to 1.4×10^−11^ kJ cell^−1^ y^−1^:

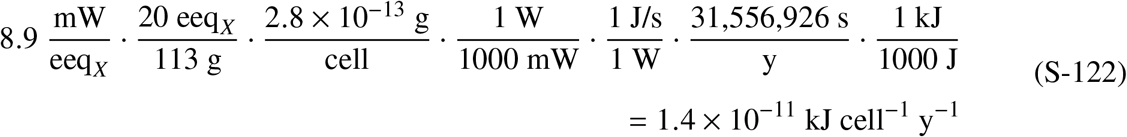

However, Lever et al. ^24^ reports a minimum energy dissipation of 1 × 10^−15^ kJ cell^−1^ y^−1^ at the extreme starvation condition in the deep sediment column, which is about 4 orders of magnitude lower than at this 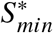. Under extreme starvation, cells break down all machinery that is not essential for bare minimum survival. Under these conditions, microorganisms persist through net decay over thousands and millions of years, gradually yielding the ground to the environmental strategy. This is markedly different from 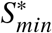, which sets a minimum energy dissipation requirement to maintain steady-state biomass and invade the environment. Microorganisms have developed strategies such as dormancy, sporulation, and other persistence mechanisms to survive unfavorable growth conditions under conditions where 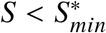.

